# Single-cell CRISPR activation screens in primary B cells discover gene regulatory mechanisms for hundreds of autoimmune risk loci

**DOI:** 10.64898/2026.03.01.708923

**Authors:** Viacheslav A. Kriachkov, Jeralyn Wen Hui Ching, James Lancaster, Davide Vespasiani, Nicholas Denny, Joseph C. Hamley, Liam Gubbels, Esther Bandala Sanchez, Melanie Neeland, Eric Levi, Karen Davies, Shivanthan Shanthikumar, Galina Shevchenko, Vanessa L. Bryant, Daniel J. Hodson, James O. J. Davies, Hamish W. King

**Affiliations:** Walter and Eliza Hall Institute of Medical Research; Parkville, VIC 3052 Australia; Department of Medical Biology, University of Melbourne; Parkville, VIC 3052 Australia; MRC Molecular Haematology Unit, MRC Weatherall Institute of Molecular Medicine, University of Oxford; Oxford OX3 9DS, United Kingdom; Infection, Immunity and Global Health Theme, Murdoch Children’s Research Institute; Parkville, VIC 3052 Australia; Otolaryngology Department, Royal Children’s Hospital; Parkville, VIC 3052 Australia; Respiratory and Sleep Medicine, Royal Children’s Hospital; Parkville, VIC 3052 Australia; Department of Paediatrics, University of Melbourne; Parkville, VIC 3052 Australia; Cambridge Stem Cell Institute; Cambridge CB2 0AW, United Kingdom; Department of Haematology, University of Cambridge; Cambridge CB2 0RE, United Kingdom; National Institute of Health Research Biomedical Research Centre, University of Oxford; Oxford OX3 9DU, United Kingdom

## Abstract

Genome-wide association studies (GWAS) have discovered thousands of genetic variants linked to autoimmune disease, and yet the molecular pathways underlying autoimmunity have remained elusive. A key challenge is that >90% of identified GWAS risk loci are in non-coding genomic regions making it difficult to predict their relevance to disease. Here, we have curated fine-mapped non-coding risk variants from over 30 different autoimmune traits including common conditions such as systemic lupus erythematosus (SLE), Crohn’s disease, and multiple sclerosis, and reveal shared genetic signatures between diverse autoimmune diseases. We subsequently performed a high-throughput single-cell multi-omic CRISPR activation screen targeting 763 autoimmune risk loci in primary human B cells (a highly relevant cell type to autoimmune diseases) and discover 524 *cis*-regulatory target gene effects for 378 risk loci, with many risk loci regulating multiple gene targets. This Single Cell Analysis of Non-coding Distal Autoimmune Loci (SCANDAL) provides a powerful experimental resource linking non-coding risk loci to many disease-relevant genes, including lowly-expressed cytokines and transcription factors for which perturbation effects can be difficult to quantify with other CRISPR-based strategies. We reveal how increased transcriptional activity at one non-coding risk locus can drive transcription at other risk loci within the same regulatory landscape that may be relevant to understand genetic pleiotropy of autoimmune diseases. Finally, we quantified allele-specific effects on target gene expression with massive parallel reporter assays and prime editing to discover a gain-of-function variant associated with SLE that controls expression of the transcription factor *REL*/cREL which subsequently binds dozens of risk loci and target genes associated with different autoimmune diseases. Our study provides a valuable resource linking non-coding risk loci with their *cis*-regulatory target genes and advances our understanding of the shared genetic networks and mechanisms involved in autoimmunity.

## Introduction

A central challenge of human genetics during the era of genome-wide association studies (GWAS) has been to move from variant discovery to biological mechanism. Over the last two decades, GWAS have identified tens of thousands of common genetic variants statistically associated with diverse physiological traits and disease risk (*1*). However, most of these genetic signals fall in non-coding regions of the genome, and it has proved difficult to interpret the potential consequences of genetic variants on target gene expression and disease mechanism (*2, 3*).

Autoimmune diseases result from a loss of immune tolerance to normally harmless endogenous or exogenous antigens, and can be caused by dysregulation in the selection, differentiation, or function of immune cells. Self-reactive T and B cell lymphocytes and the production of autoantibodies are common features of many autoimmune traits. These diseases have a strong heritable component, as evidenced by familial and twin studies (*4, 5*), and apart from rare monogenic cases (*6, 7*), most autoimmune disorders are considered prototypical complex genetic diseases with common single nucleotide polymorphisms (SNPs) collectively mediating lifetime risk of developing one or more autoimmune disease. Furthermore, polygenic risk scores and genetic correlation analyses indicate broad pleiotropy and show that shared risk loci converge on key immunological pathways such as lymphocyte activation, cytokine signalling, and innate immune regulation (*8–10*). The mechanisms by which these genetic risk loci modulate these phenotypes remains a major unsolved challenge in the field.

Cell type-resolved functional genomics such as chromatin accessibility, histone modifications, chromosome confirmation capture (3C)-based assays and gene expression have consistently shown enrichment of GWAS signals at *cis*-regulatory elements (CREs) across diverse immune cell types and states such as macrophages, T cells, and B cells (*11–15*). Targeting putative CREs with catalytically dead CRISPR/Cas9 (dCas9) fused with inhibitory (CRISPRi) or activating (CRISPRa) domains can modulate transcription of gene promoters dependent upon those loci for normal expression (*16, 17*). These strategies have allowed for identification of *cis*-regulatory gene targets of genomic regions containing non-coding GWAS variants in immortalised cancer cell lines (*17–20*) or more recently primary T cells (*21, 22*). However, other disease-relevant cell types such as B cells have proved refractory to viral delivery strategies required for high-throughput functional genomic screens. Similarly, quantification of variant-level effects on gene expression with massively parallel reporter assays (MPRA) (*22, 23*), and more recently *in situ* base- and prime-editing (*17, 24*), have yet to be applied widely in diverse primary cell types.

Here, we investigate how non-coding risk loci regulate expression levels of their target genes to modulate autoimmune disease risk. We identify a shared genetic architecture across diverse autoimmune diseases and systematically map *cis*-regulatory target genes of hundreds of non-coding risk loci by performing a highly-multiplexed single-cell multi-omic CRISPR activation screen in primary human B cells, a disease-relevant cell type that is traditionally refractory to high-throughput genomic screens. This Single Cell Analysis of Non-coding Distal Autoimmune Loci (SCANDAL) allowed us to uncover diverse and complex *cis*-regulatory effects by autoimmune risk loci in controlling expression of genes involved with disease-relevant phenotypes, including gene targets with low expression levels that would be missed by CRISPRi screening approaches. Finally, we employ MPRA and *in situ* prime editing to quantify loss- and gain-of-function effects of non-coding variants. Together, our findings represent a step toward systematically linking non-coding risk variants to disease-relevant mechanisms within the human immune system.

## Results

### A meta-analysis of the shared genetic architecture for autoimmune disease risk

We curated 5,655 statistically fine-mapped risk variants for 31 autoimmune GWAS traits from OpenTargets Genetics and Probabilistic Identification of Causal SNPs (PICS2) (*25, 26*) (table S1). This pan-autoimmune resource includes common traits such as multiple sclerosis (MS), systemic lupus erythematosus (SLE), inflammatory bowel disease (IBD) and rheumatoid arthritis (RA), as well as less common diseases like Sjögren’s disease, systemic sclerosis and Graves’ disease (**Fig. 1A**), although the available genetic data per trait did not always correlate with its prevalence (*27*) (fig. S1A). Over 40% of risk variants had higher frequencies in non-European populations that are often under-represented in genetic studies (**Fig. 1B**). While the Human Leukocyte Antigen (HLA) locus on chromosome 6 is a major genetic risk signal for autoimmunity, over 93% of risk variants in our pan-autoimmune resource were non-HLA loci (**Fig. 1C**, fig. S1B).

**Fig. 1.**
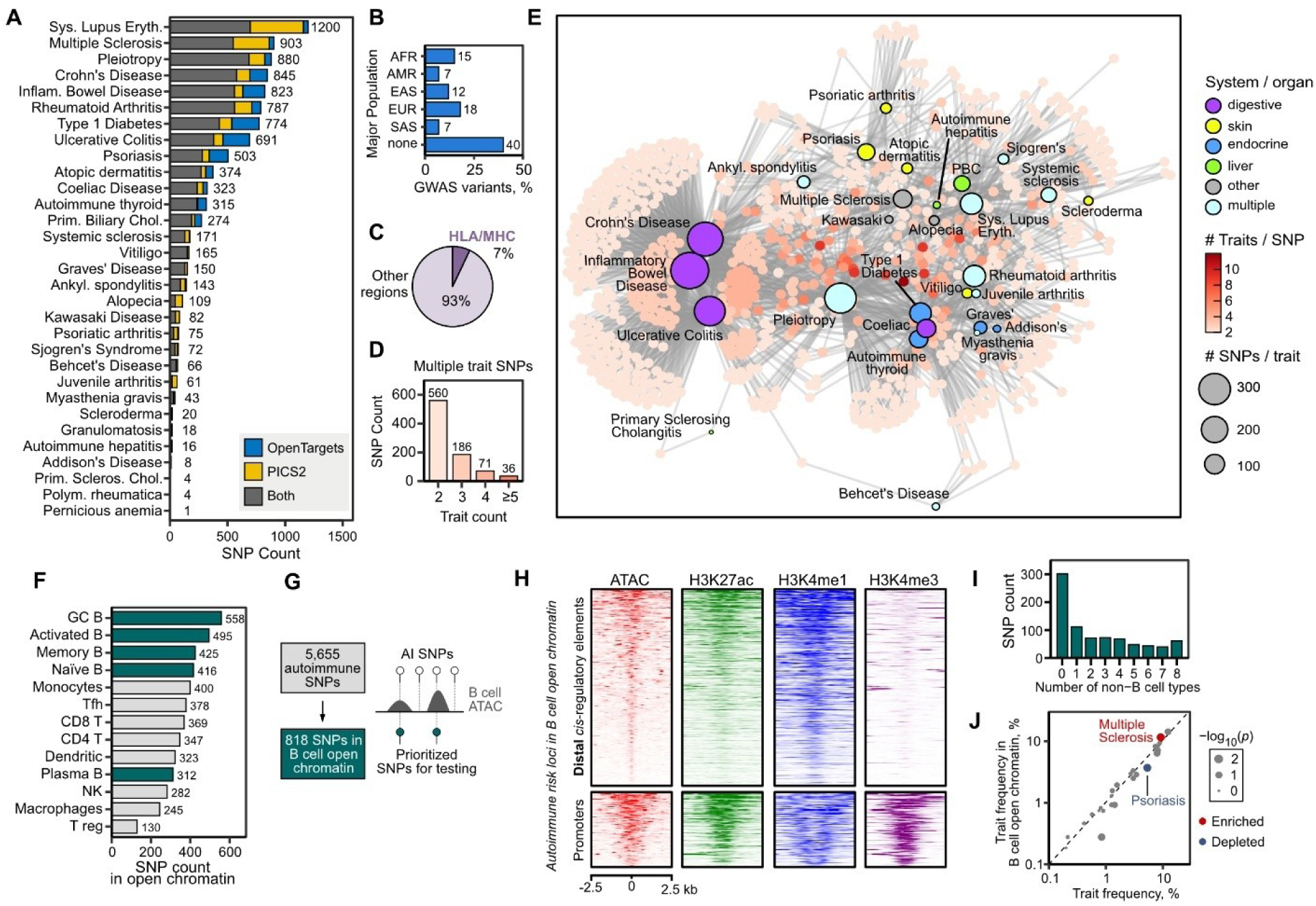
A pan-autoimmune disease resource of fine-mapped GWAS risk variants. (**A**) Trait frequency for 5,655 autoimmune risk SNPs from PICS2 and OpenTargets Genetics. (**B**) Predominant population of SNPs across different ancestry groups (AFR = African, AMR = admixed American, EAS = East Asian, EUR = European, SAS = South-East Asian). (**C**) Percentage of SNPs located within HLA/MHC cluster. (**D**) Trait frequency for 853 multi-trait SNPs. (**E**) Shared trait associations for multi-trait SNPs visualised as a force-directed graph. (**F**) Chromatin accessibility at autoimmune SNPs in immune cells. (**G**) Prioritization of 818 autoimmune SNPs based on chromatin accessibility in B cells. (**H**) GC B cells chromatin accessibility (ATAC) and histone marks at 818 autoimmune SNPs centered on the middle of open chromatin peak. (**I**) Chromatin accessibility of 818 SNPs from B cell open chromatin in non-B cell types. (**J**) Trait enrichment after functional prioritisation with B cell open chromatin. Red or blue denotes significant increase or decrease by Fisher exact test.

We identified 853 variants (15.1%) linked with two or more autoimmune traits (**Fig. 1D**), and network graph analysis of these multi-trait loci revealed a shared genetic architecture that broadly separated gut-associated inflammatory disease traits (Crohn’s disease, ulcerative colitis) from more systemic autoimmune diseases, with related traits often clustering together (*e.g.* psoriatic arthritis, psoriasis, atopic dermatitis) and pleiotropic GWAS traits clustering in the centre (**Fig. 1E**). Intriguingly, coeliac disease had greater degree of shared risk variants with systemic and non-gut-related autoimmune traits, perhaps consistent with importance of gluten-reactive T lymphocyte responses in this disease compared to other inflammatory gut diseases (*28*). Notably, the vast majority (90.5%) of variants in our pan-autoimmune disease resource were in non-coding genomic regions with an average distance of 77.4 kb to the nearest transcription start site (TSS) (fig. S1C-D), making it challenging to interpret their relevance or role in disease.

### Prioritizing autoimmune risk loci with regulatory potential in human B cells

We next used single-cell chromatin accessibility maps from diverse immune cell subsets spanning bone marrow, peripheral blood and secondary lymphoid organs (*14*) to prioritize potentially causal variants that mediate disease risk through changes in gene expression. This identified 1013 variants in open chromatin in at least one immune cell type, with germinal centre (GC) B cells exhibiting the most variants within accessible chromatin regions compared to other examined cell types (**Fig. 1F**). Given that self-reactive B cells and B cell-derived autoantibodies are a common feature in many autoimmune diseases, and that the GC is a major site where autoreactive B cells must be controlled to prevent autoimmunity (*29, 30*), we prioritised 818 risk variants overlapping 684 putative *cis*-regulatory elements (CRE) in GC and other B cell subsets (**Fig. 1G**, table S2) with the majority enriched for chromatin modifications consistent with distal CREs such as high H3K27ac, high H3K4me1 and low H3K4me3 (**Fig. 1H**, fig. S1E-F). Many (57%) of these loci exhibited chromatin accessibility in diverse immune cell types (**Fig. 1I**), suggesting broader potential relevance beyond lineage-specific mechanisms, and while our prioritisation of risk variants did not enrich individual traits except for MS (**Fig. 1J**, fig. S1G), it did significantly enrich for risk variants associated with multiple autoimmune traits (19.2% vs 14.6%; *p*<0.00082). However, it remained difficult to infer how, if at all, these non-coding loci regulate gene expression and cellular phenotypes to mediate increased risk for autoimmune disease.

### Tripartite delivery of CRISPR-Synergistic Activation Mediator in primary B cells

We therefore set out to experimentally map the *cis*-regulatory target genes of these non-coding risk loci in primary B cells. However, B cells are refractory to conventional VSV-G pseudotyped viral transduction and high-throughput functional genomic screens in B cells have remained extremely challenging. We engineered a CRISPR-Synergistic Activation Mediator (CRISPR-SAM) system (**Fig. 2A**) (*31*) for viral delivery into primary human tonsillar GC B cells with RD114-type retroviruses encoding dCas9-VP64 and the synergistic activator MS2-coat protein (MCP)-p65-HSF1, along with GaLV-type lentivirus encoding sgRNA fused to MS2 loops to target CRISPR-SAM to specific loci (**Fig. 2B-C**, fig. S2A-D). This allowed CRISPR activation of targeted genes in primary B cells without additional modification (<5-10 days in culture) or primary B cells that over-express BCL6/BCL2 to limit plasma cell differentiation and prolong survival in culture (10+ days) (fig. S2E). To explore the dynamic range of CRISPR-SAM activation in primary B cells, we targeted either the complete CRISPR-SAM (dCas9-VP64 + MCP-p65-HSF1) or only dCas9-VP64 to either silent (*CD5*), moderately- (*CD86*) or highly-expressed (*CD19*) gene promoters, and found that CRISPR-SAM consistently resulted in higher activation than dCas9-VP64 alone at the tested loci (**Fig. 2D**, fig. S2F-G).

**Fig. 2.**
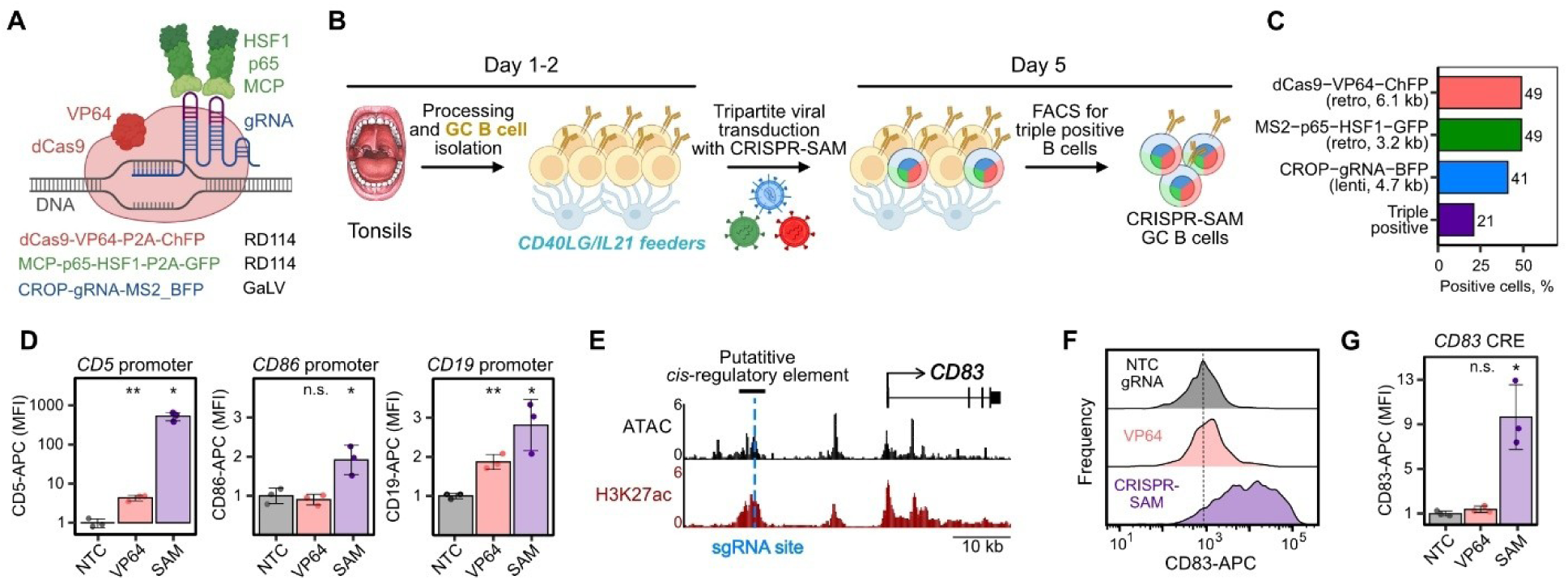
CRISPR-Synergistic Activation Mediator (CRISPR-SAM) in primary human B cells to modulate regulatory activity of autoimmune risk loci. (**A**) Schematic of CRISPR-SAM activator, comprising catalytically dead Cas9 (dCas9) fused with VP64, activators p65 and HSF1 fused to MS2 coat protein (MCP) that binds MS2 loops in the sgRNA structure. Viral delivery strategy (RD114 retrovirus, GaLV lentivirus) is detailed for each construct. (**B**) Timeline for CRISPR-SAM activation in human GC B cells. (**C**) Transduction efficiencies for individual and triple delivery of CRISPR-SAM components. (**D**) Mean fluorescence intensities (MFI) for CD5, CD86, or CD19 after transduction of GC B cells with promoter-targeting sgRNA for each gene and dCas9-VP64 only or dCas9-VP64+MS2-p65-HSF1 (CRISPR-SAM). MFIs are normalised to samples with non-targeting control (NTC) sgRNA and CRISPR-SAM (n=3, ±SD). (**E**) Genome snapshot of CRISPR-SAM sgRNA-targeting putative *cis*-regulatory element (CRE) upstream of *CD83*. GC B cell ATAC-seq and H3K27ac ChIP-seq are shown. (**F**) Representative histogram of CD83 surface protein expression after CRISPR-SAM targeting of *CD83* putative CRE in GC B cells. Dashed line shows NTC MFI. (**G**) CD83 MFI values relative to NTC (n=3, ±SD) after CRISPR-SAM at *CD83* CRE. *p*-values from Student’s t-test are shown (* < 0.05; ** < 0.01).

Finally, to test whether this approach could be used to map *cis*-regulatory target genes of non-coding loci, we targeted CRISPR-SAM to a putatitive CRE located 22 kb upstream of the *CD83* promoter (**Fig. 2E**). This significantly up-regulated *CD83* (**Fig. 2F-G**) and is consistent with enhancer-promoter co-accessibility (*14*), expression quantitative trait loci (eQTL) (*32*), and CRISPRi FLOW-FISH (*18*) analyses supporting *cis*-regulatory control of the *CD83* promoter by the targeted region. This therefore demonstrated the utility of CRISPR-SAM to map *cis*-regulatory targets of non-coding risk loci in primary human B cells.

### Single-Cell Analysis of Non-coding Distal Autoimmune Loci (SCANDAL)

We next generated a sgRNA library to target CRISPR-SAM to 763 CREs encoding 791 fine-mapped autoimmune risk variants so that we could experimentally map their *cis*-regulatory target genes (**Fig. 3A**, table S3). We designed three sgRNAs per locus as well as control sgRNAs to target 30 inert genomic regions at either gene deserts (no annotated genes ±500 kb) or in closed chromatin in neighbouring proximity (<10kb) to TSSs, in addition to 18 non-targeting (NT) sgRNAs (**Fig. 3A**, fig. S3A-B). Tonsil-derived GC B cells were transduced with CRISPR-SAM components and the sgRNA library and labelled with oligonucleotide-tagged antibodies for 137 cell surface proteins before single-cell capture and library generation. This **S**ingle **C**ell **A**nalysis of **N**on-coding **D**istal **A**utoimmune Loci (SCANDAL) could then be used to empirically test the regulatory potential for hundreds of risk loci in a highly multiplexed experiment. We analysed 201,191 cells with at least one sgRNA (MOI=1.38) and an average of 116 cells per sgRNA (average of 347 cells per locus) after doublet removal and stringent quality control (**Fig. 3B**, fig. S3C-D). To identify *cis*-regulatory targets of autoimmune risk loci, we tested for differential expression of all protein-coding genes or antibody-derived tags (ADT) encoded within 500kb of each locus (**Fig. 3A**, fig. S3E-F) (*17, 33*). To reduce the risk of false positives, we set an additional requirement that at least two out of the three sgRNAs had to independently return significant and concordant effects (fig. S3G). Importantly, we observed no significant effects on nearby gene expression for sgRNAs targeting the CRISPR-SAM system to closed chromatin in gene deserts or near gene promoters (**Fig. 3C**). Our SCANDAL screen of non-coding risk loci identified ≥1 significant *cis-*regulatory target genes for 378 out of 763 targeted loci (49.5%) (**Fig. 3C-D**), such as for the autoimmune thyroid disease-associated rs7441808 that specifically regulated expression of *RBPJ* (required to prevent spontaneous GC formation in mice (*34*)) over a genomic distance of 180 kb (**Fig. 3E-F**) (*22*). In total we mapped 524 significant locus-gene pairs (**Fig. 3D**, table S4-5), the majority of which had statistical support from published eQTL studies (**Fig. 3G**). We identified 10 or more *cis*-regulatory target genes for 14 different autoimmune traits (**Fig. 3H**) and we did not observe enrichment of any specific autoimmune traits amongst the loci with *cis*-regulatory targets (fig. S4A-B). About a third of identified target genes (127/362; 35%) were mapped to more than one trait and these multi-trait genes shared similar patterns between traits to our previous multi-trait variant analysis (**Fig. 1E**, fig. S4C). This supports that SCANDAL *cis*-regulatory target genes mediate some of the shared genetic architecture of autoimmunity.

**Fig. 3.**
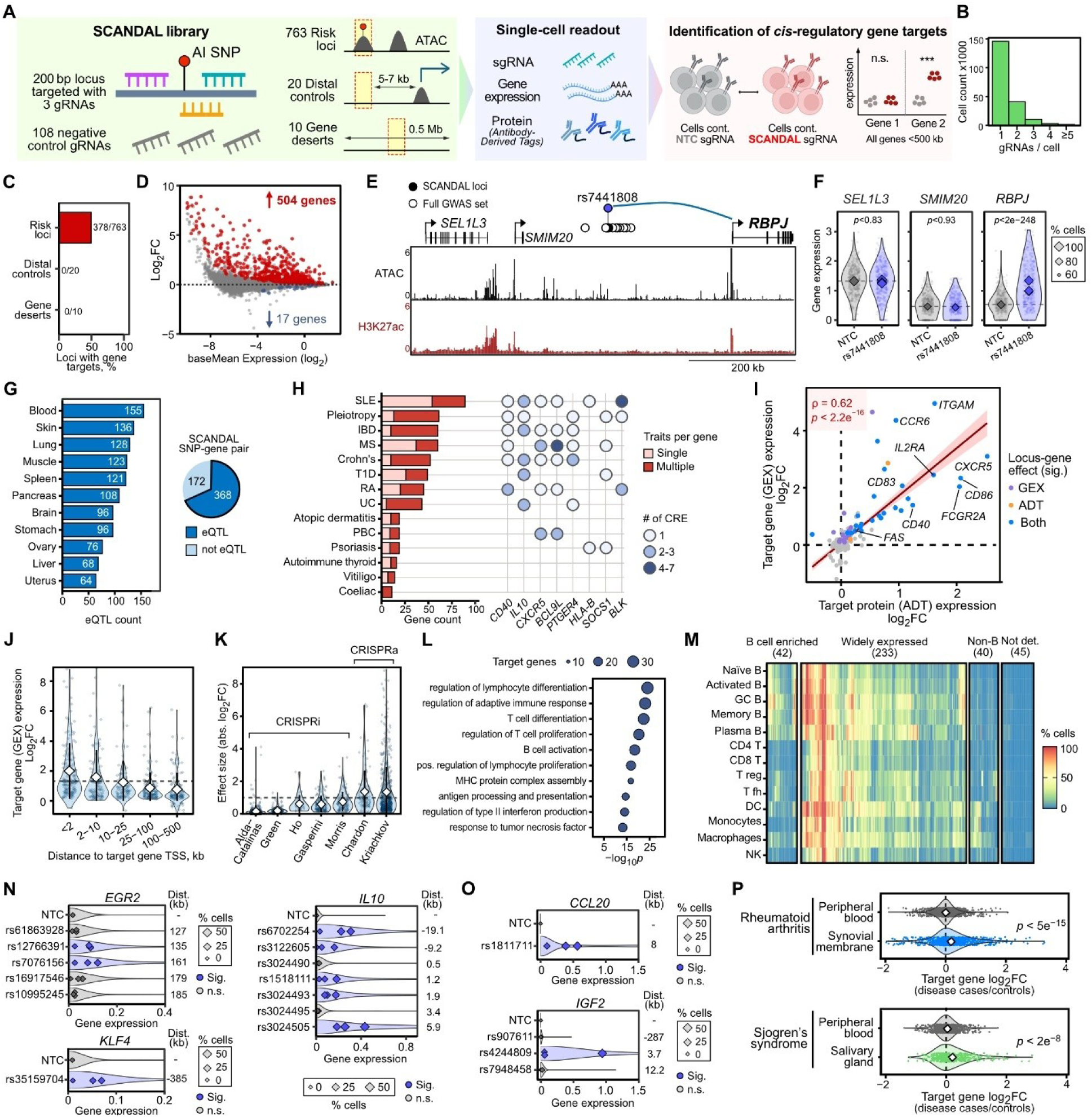
Single-cell CRISPR activation screens in human B cells identifies hundreds of gene targets regulated by autoimmune risk loci. (**A**) Schematic of SCANDAL sgRNA library design for autoimmune risk loci and control regions, single-cell capture and identification of *cis*-regulatory targets in response to CRISPR-SAM perturbation. (**B**) Multiplicity-of-infection (MOI) of SCANDAL library in GC B cells. (**C**) Relative frequency of tested loci with significant *cis*-regulatory effects on gene expression within 500 kb. (**D**) Differential expression of genes within 500 kb for all targeted SCANDAL loci. Significant locus-gene effects are denoted in red and blue. (**E**) Genome snapshot of CRISPR-SAM targeting at rs7441808, including *cis*-regulatory effect on target gene *RPBJ* shown as curved line. (**F**) Expression levels of genes in (E) for cells containing rs7441808-targeting or non-targeting control (NTC) sgRNAs. Points reflect single cell expression values, diamonds indicate mean expression values, size corresponds to cell percentage the gene was detected, and Benjamini-Hochberg-adjusted SCEPTRE *p*-values are shown. (**G**) Number of SCANDAL locus-gene pairs with GTEX eQTL support in different tissues. (**H**) Number of SCANDAL target genes mapped per trait for top 14 autoimmune traits, classified as to whether they were targets for loci linked with multiple traits. Gene-trait pairs for eight target genes with the most traits, including the number of independent SCANDAL *cis*-regulatory effects underlying the associations. (**I**) Comparison between gene expression (GEX) and protein levels (antibody derived tags; ADT) *cis*-regulatory effects for SCANDAL loci. Spearman’s rank correlation (ρ) and *p*-value are shown. (**J**) Changes in SCANDAL targets by genomic distance to risk locus. (**K**) Absolute perturbation effect sizes for *cis*-regulatory targets from different studies. (**L**) Gene ontology for 362 SCANDAL target genes. (**M**) Expression of SCANDAL targets across immune cell types. (**N-O**) SCANDAL effects of either lowly expressed (**N**) or not detected at baseline (**O**) *cis*-regulatory target genes. (**P**) Change in SCANDAL target gene expression between disease and controls in peripheral blood and disease-relevant tissues. *p*-values from Student’s t-test are shown.

Many of the *cis*-regulatory effects we observed on transcription also led to changes in protein expression, with a high degree of concordance between gene expression and cell surface protein changes after CRISPR-SAM activation (table S6), including important immune-related genes such as *CD40*, *FAS* (CD95), *IL2RA* (CD25), *CD83* and *CXCR5* (**Fig. 3I**, fig. S5A-F). Furthermore, SCANDAL allowed robust identification of *cis*-regulatory relationships with an average effect size of 2.5-fold on target gene expression, and this was only partially dependent on proximity to the target TSS with many examples of large effect sizes (e.g. 2- to 8-fold) over hundreds of kilobases (**Fig. 3J**). These effect sizes were consistently higher than those observed in CRISPRi-based perturbation screens of non-coding risk loci (**Fig. 3K**), suggesting that CRISPRa-based screens like SCANDAL have a higher dynamic range to capture *cis*-regulatory relationships of non-coding risk loci, including in challenging cell types such as primary B cells.

### SCANDAL *cis*-regulatory targets include lowly expressed and disease-relevant genes

We identified 362 unique *cis*-regulatory target genes whose expression was regulated by at least one autoimmune risk locus, and these SCANDAL target genes were strongly enriched for immune-related pathways (**Fig. 3L**) and widely expressed by different immune cell types (**Fig. 3M**). We also observed that SCANDAL could map lowly expressed *cis*-regulatory target genes like transcription factors (*EGR2*, *KLF4*), cytokines (*IL10*, *IL27*, *IL12A*), and immune-modulatory genes such as *CCR6*/CD196 (chemokine receptor required for B cell chemotactic response) (**Fig. 3N**, fig. S6A-C). SCANDAL also enabled us to map *cis*-regulatory target genes normally expressed by cell types other than B cells such as *CCL20* (ligand for CCR6; normally expressed by monocytes and T cells) and placenta-enriched insulin-growth factor 2 (IGF2) that was regulated by a locus encoding the type 1 diabetes-associated rs4244809 (**Fig. 3O**). Furthermore, when we examined the differential expression of SCANDAL target genes in autoimmune disease cohorts relative to healthy controls, they were more likely to be differentially expressed only in disease-relevant tissues, and not in peripheral blood, such as synovium in rheumatoid arthritis or salivary gland in Sjögren’s disease (**Fig. 3P**, fig. S6D). These findings highlight how SCANDAL can probe the regulatory potential of non-coding risk loci to reveal biologically- and disease-relevant *cis*-regulatory target genes.

### SCANDAL maps *cis*-regulatory targets over large genomic distances, including non-proximal genes

The majority of risk loci with gene targets from SCANDAL (244/378; 64.6%) were in distal (non-promoter) genomic regions (**Fig. 4A**). More than a quarter of *cis*-regulatory gene targets of these distal loci were greater than 100 kb away from their risk locus, highlighting the prevalence of long-distance gene regulatory control of target genes by non-coding risk loci. We next performed Micro-Capture-C (MCC) (*35, 36*) for selected SCANDAL loci or target gene promoters to determine whether these were consistent with a physical interaction between the risk loci and identified SCANDAL target genes. At the *ID2* locus, our SCANDAL screen identified a CRISPR-SAM-responsive CRE (containing the atopic dermatitis-associated SNP rs10174949) that regulates expression of *ID2* located 380 kb upstream (**Fig. 4B-C**). *ID2* encodes a transcriptional regulator with reported roles in B cell differentiation and class switch recombination (*37, 38*) and has been implicated in Th17-mediated autoimmunity (*39*). Analysis at this locus with MCC showed a distinct signal at *ID2* promoter, supporting a direct role of this risk locus in regulating *ID2* expression (**Fig. 4B**). Similarly, we found five non-coding regulatory elements in an apparent locus-control region 189-286 kb upstream of *PTGER4* (fig. S7A-B). Although these *cis*-regulatory effects occurred over large genomic distances (>100 kb), in both cases the identified target gene was the gene nearest to the tested risk locus. Overall however, the distance between experimentally-mapped SCANDAL target genes and their risk locus was significantly greater compared to the distance to most proximal TSS for autoimmune risk loci (**Fig. 4D**; *p*<3.2×10^-14^). In fact, 159/378 risk loci regulated a gene target that was not the nearest gene (**Fig. 4E**). Indeed, SCANDAL identified 58 regulatory elements that skipped the nearest gene to only regulate more distal targets (fig. S7C-G), as in the case of rs67289879 (Crohn’s disease, IBD) locus which regulated *CCND3 (98 kb* away), but not *TAF8* (11 kb away) (**Fig. 4F-G**). Other examples of CREs with a specific effect on a non-adjacent gene include rs10152590 (RA) locus that regulated *TLE3* (>180 kb; fig. S7C-D) and rs78037977 (T1D, vitiligo) locus that controlled *TNFSF4* over 460 kb away (fig. S7E-F) with no changes in other nearby genes. These observations demonstrate that SCANDAL can specifically and accurately detect *cis*-regulatory relationships over a wide-range of genomic distances and complex regulatory landscapes.

**Fig. 4.**
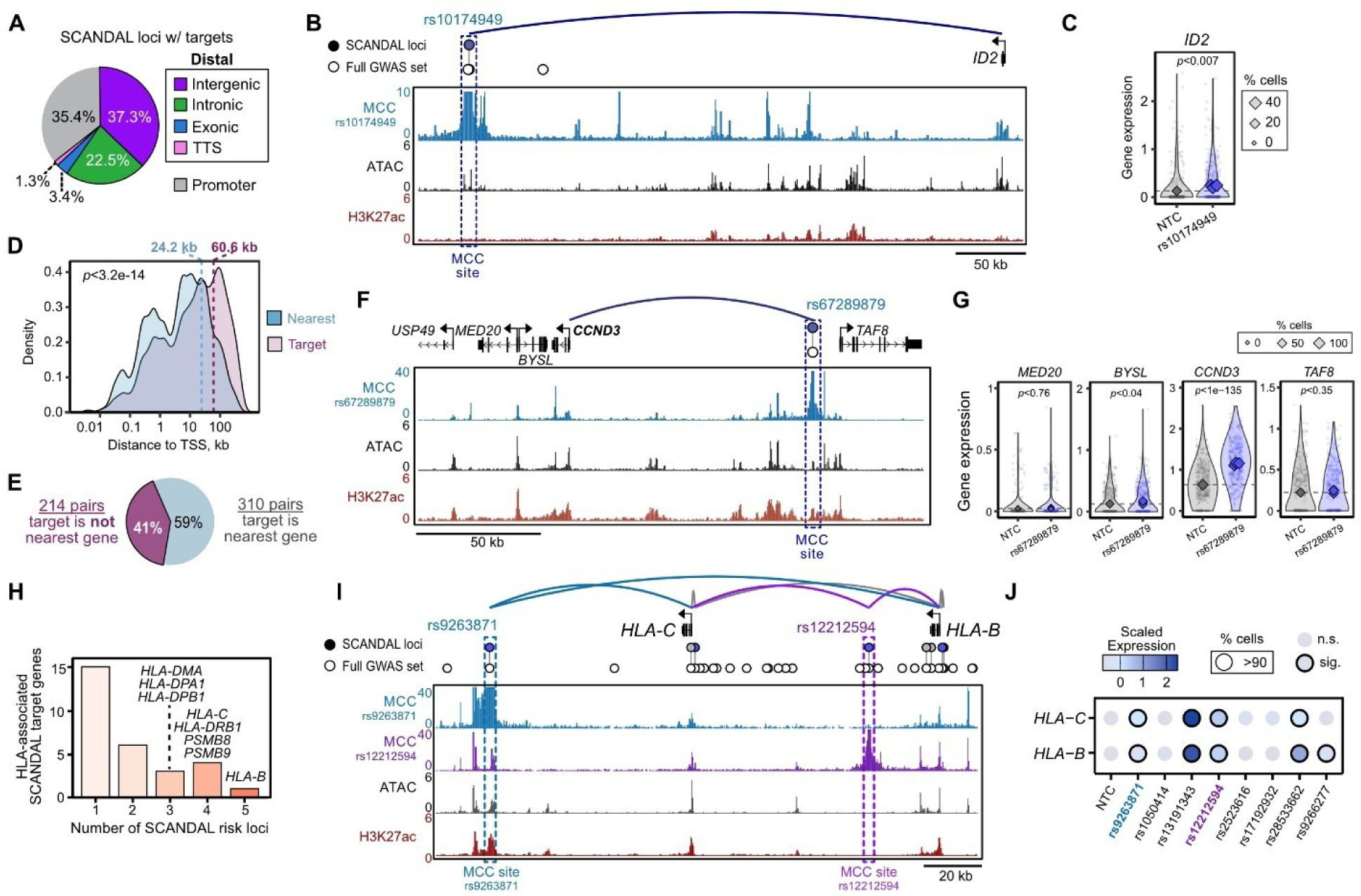
Distal regulatory gene targets of risk loci identified by SCANDAL and micro-Capture-C. (**A**) Genomic context of SCANDAL risk loci (378) with mapped *cis-*regulatory target genes. TTS = transcription termination site. (**B**) Genome snapshot of CRISPR-SAM targeting and Micro-Capture-C (MCC) for rs10174949, including SCANDAL *cis*-regulatory effect on target gene *ID2* highlighted as curved line. (**C**) Expression levels of *ID2* for cells containing rs10174949-targeting or non-targeting control sgRNAs in SCANDAL. Circular points reflect single cell expression values, while diamonds indicate mean expression values for all cells containing individual sgRNAs where size corresponds to percentage of cells in which the gene was detected. *p* denotes Benjamini-Hochberg-adjusted SCEPTRE *p*-value. (**D**) Distance from SCANDAL risk loci to either nearest TSS vs TSS of experimentally-mapped target gene. Dashed lines show the median distance. *p*-value from Student’s t-test is shown. (**E**) Frequency that gene target for a risk locus is the nearest gene or not. (**F**) Genome snapshot of CRISPR-SAM targeting and Micro-Capture-C (MCC) for rs67289879, including SCANDAL *cis*-regulatory effect on target gene *CCND3* highlighted as curved line. (**G**) Expression of *MED20*, *BYSL*, *CCND3* and *TAF8* in cells containing rs67289879-targeting or non-targeting control sgRNAs in SCANDAL. (**H**) Frequency of SCANDAL target genes within the HLA super-locus (chr6:28510120-33480577), including the number of independent risk loci for each gene. (**I**) Genome snapshot of SCANDAL targeting of 8 loci and Micro-Capture-C (MCC) for rs9263871 and rs12212594 near *HLA-C* and *HLA-B*. (**J**) Scaled expression (z-scores) of *HLA-C* and *HLA-B* for cells containing sgRNAs targeting 8 different autoimmune risk loci or non-targeting control sgRNAs (NTC) in SCANDAL.

### Non-coding regulatory elements that modulate MHC gene expression

The HLA locus encodes the Major Histocompatibility Complex (MHC) proteins and co-factors and has the strongest known genetic associations with autoimmunity, including protein-coding variation in HLA alleles (*40, 41*). We targeted 87 risk loci at the HLA super-locus (chr6:28510120-33480577) with CRISPR-SAM to determine their *cis*-regulatory potential and identified gene targets for 38 SCANDAL loci (44%), including MHC class I genes (*HLA-A*, *HLA-B*, *HLA-C*), MHC class II genes (*HLA-DRB1*, *HLA-DRA*, *HLA-DRB5*), and associated genes involved with peptide processing and presentation (*PSMB8*, *PSMB9*, *TAP1*, *TAP2*, *MICA*) (**Fig. 4H**). SCANDAL identified CREs that regulated expression of single HLA genes (*e.g.* regions encoding rs34941730 and rs6935053 for HLA-A) (fig. S7H-I). Intriguingly, we also found risk loci that regulated multiple HLA genes (*e.g.* HLA-B and HLA-C by regions encoding rs9263871 and rs12212594) (**Fig. 4I-J**, fig. S7J), with physical interaction support from MCC confirming the specificity of these functional effects. These observations highlight that in addition to coding HLA variants, non-coding risk loci reflect an under-appreciated risk mechanism to modulate MHC levels in autoimmunity (*42, 43*).

### Transcriptional activity of SCANDAL *cis-*regulatory elements

By targeting CRISPR-SAM activation machinery to hundreds of CREs throughout the genome, SCANDAL provided a unique opportunity to investigate general principles of *cis*-regulatory landscapes. We found that distal CREs with high chromatin accessibility and H3K4me1 levels were more likely to have a *cis*-regulatory target gene, with an unexpectedly modest difference for H3K27ac (**Fig. 5A**). Given that RNA polymerase II-dependent transcription of short and unstable RNAs from CREs is linked with regulation of their target genes (*44*), we quantified transcribed CRE (tCRE) activity at autoimmune risk loci using Single-Cell Analysis of Five-prime Ends (SCAFE) to measure 5’-derived TSS RNAs (**Fig. 5B**, table S7) (*45*). We observed no relationship between baseline tCRE activity and the probability they had a *cis*-regulatory target (**Fig. 5C**), although detection sensitivity of the single-cell RNA-seq readout likely limited our ability to detect low expression of tCRE-derived RNAs. We therefore considered whether the use of CRISPR activation at SCANDAL loci would drive tCRE above the detection threshold and allow us to investigate their properties, analogous to our observations for lowly expressed mRNA transcripts (**Fig. 3N-O**). Indeed, 69% (375/547) of distal CREs showed detectable tCRE activity after CRISPR-SAM activation compared to 36% at baseline (table S8) and, notably, we found that distal loci with increased tCRE activity were more likely to have a *cis*-regulatory gene target (**Fig. 5D**). Importantly, we detected negligible or no tCRE signal at control genomic loci after CRISPR-SAM targeting (**Fig. 5D**), supporting that CRISPR-SAM itself is not sufficient to recruit active RNA polymerase II-dependent transcription in the absence of pre-existing regulatory activity. Single-cell differential expression analysis revealed 168 distal loci with significant increases in tCRE activity after CRISPR-SAM (**Fig. 5E**, table S8) and we found that this CRISPR-SAM-dependent increase in tCRE activity correlated with the change in SCANDAL target gene expression (**Fig. 5F**). Some examples of CRISPR-SAM-dependent tCRE activity include rs10152590 (RA) that regulates *TLE3* (**Fig. 5G**, fig. S7C-D) and rs1874252 (SLE) that regulates *RASGRP1* (fig. S8A). These results demonstrate that CRISPR-SAM-dependent tCRE activity may provide a new approach to functional prioritization of non-coding risk loci in future studies.

**Fig. 5.**
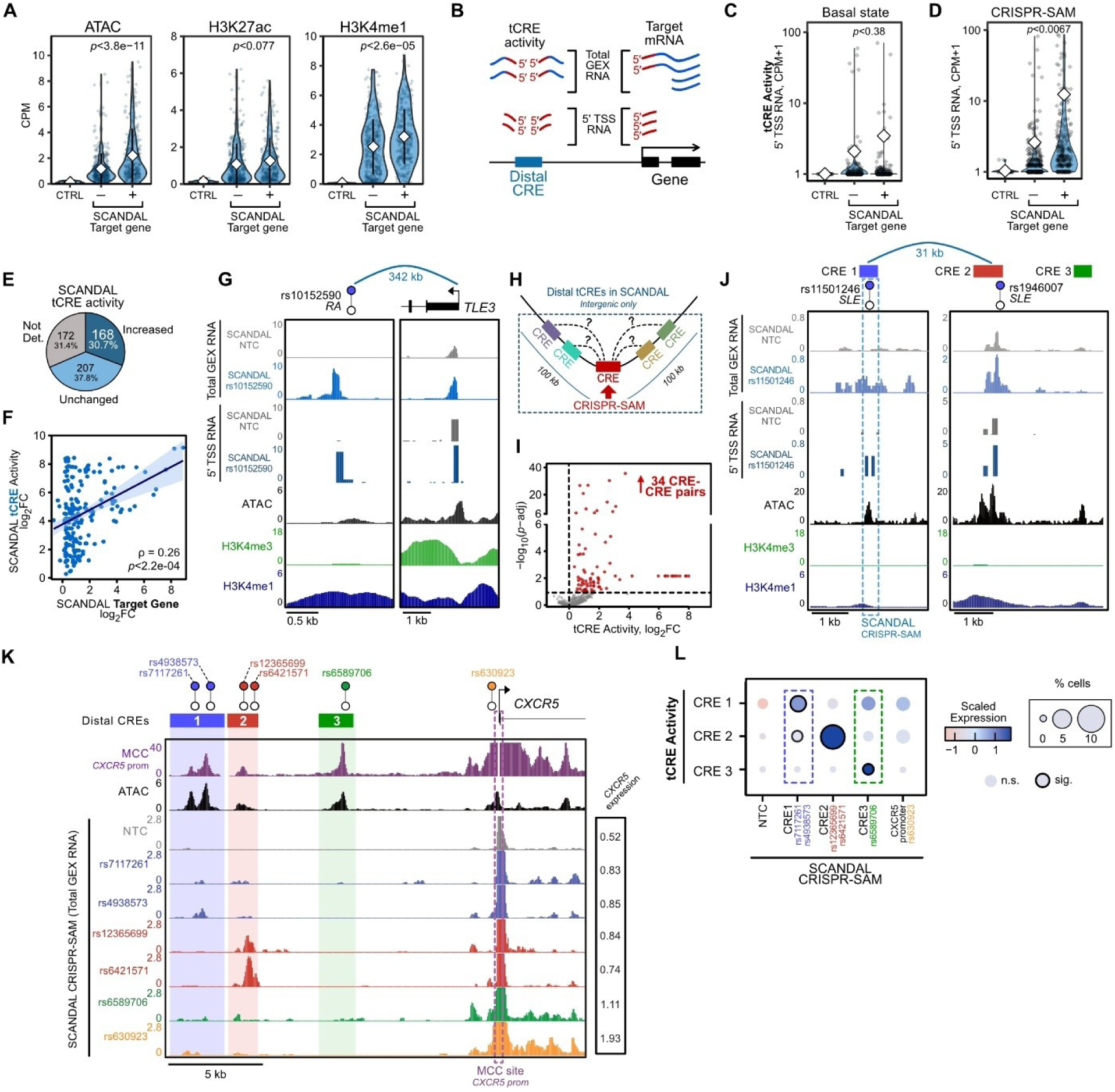
Transcriptional activity and co-regulation of non-coding autoimmune risk loci. (**A**) Chromatin accessibility and histone modification levels (CPM; counts per million) at distal SCANDAL loci (±250 bp from SNP) with or without a mapped *cis*-regulatory target gene. CTRL denotes gene desert and distal control loci. *p*-values denote Student’s t-test. (**B**) Quantification of transcriptional activity at CREs (tCRE) using total 5’ GEX RNA counts or 5’ TSS RNA counts. (**C-D**) 5’ TSS RNA counts at distal SCANDAL loci before (**C**) and after (**D**) CRISPR-SAM targeting, separated by whether a target gene was identified. *p*-values denote Student’s t-test. (**E**) Relative frequencies of distal SCANDAL loci with and without significant change in tCRE activity (total GEX RNA) after CRISPR-SAM. (**F**) CRISPR-SAM-dependent tCRE activity and target gene expression. Spearman’s rank correlation (ρ) and *p*-value are shown. (**G**) Genome snapshots of CRISPR-SAM perturbation for rs10152590 and TSS of target *TLE3*, including tCRE activity for cells containing non-targeting control sgRNAs and cells containing sgRNAs targeting rs10152590. (**H**) Differential testing for SCANDAL CRE-CRE perturbation effects following CRISPR-SAM activation of distal CRE elements. (**I**) Differential tCRE activity (total GEX RNA) after CRISPR-SAM perturbation of distal intergenic SCANDAL risk loci. Significant effects determined by SCEPTRE shown in red. (**J**) Genome snapshots of CRISPR-SAM perturbation for tCRE activity for cells containing non-targeting control sgRNAs and cells containing sgRNAs targeting rs11501246. (**K**) Genome snapshot of CRISPR-SAM perturbation and Micro-Capture-C (MCC) at the *CXCR5* locus, including total GEX RNA counts for cells containing non-targeting control sgRNAs and cells containing sgRNAs targeting 6 SCANDAL loci. Normalised expression of *CXCR5* for each unique CRISPR-SAM perturbation is reported. (**L**) Scaled expression (z-scores) for tCREs at *CXCR5* locus depicted in (K) for cells containing sgRNAs targeting CRE1-3, the *CXCR5* promoter, and non-targeting sgRNAs.

### Co-regulatory landscapes of non-coding autoimmune risk loci activity

Many *cis*-regulatory landscapes can be controlled by multiple distal CREs, although the potential for CREs to work cooperatively to regulate their target gene expression is not yet well understood (*46, 47*). We therefore sought to test for any instances of CRE-CRE co-regulation following CRISPR-SAM activation by quantifying tCRE activity at distal intergenic CREs within 100 kb of SCANDAL loci (**Fig. 5H**, fig. S8B). Surprisingly, we found 34 CRE-CRE co-regulatory relationships where up-regulation of a SCANDAL risk locus subsequently led to a significant increase in the transcriptional activity of another nearby intergenic CRE (**Fig. 5I**, table S9). Importantly, in most cases (30/34, 88%) the co-regulated CRE was not the most proximal, supporting a specificity in the observed transcriptional regulation between CREs. We observed this CRE-CRE co-regulation for 30.9% (26/84) of intergenic SCANDAL loci that themselves showed increased tCRE activity. Intriguingly, 23/26 SCANDAL loci (88%) with CRE-CRE coregulatory targets also had a mapped *cis*-regulatory gene target potentially implicating these CRE-CRE relationships in regulation of gene expression. Interestingly, we identified four examples of SCANDAL loci that were regulated by another SCANDAL risk locus, including where CRISPR-SAM activation at the rs1946007 (SLE) locus lead to transcriptional activation of a CRE encoding rs11501246 (SLE) over 30 kb away (**Fig. 5J**, fig. S8C). Both loci regulate expression of the transcription factor *ETS1* (fig. S8D), which is required to limit autoantibody production by B cells in mice (*48*). To further explore this CRE-CRE co-regulation between autoimmune risk loci, we examined a cluster of 3 tCREs upstream of *CXCR5* (**Fig. 5K-L**). While CRISPR-SAM targeting of CRE2 (rs12365699, rs6421571) led to highly specific transcriptional activity limited to itself, CRISPR-SAM targeting CRE1 and CRE3 resulted in co-activation of other CREs (**Fig. 5K-L**). MCC analysis confirmed highly specific interactions between all CREs and the *CXCR5* promoter, suggesting cooperative regulation of *CXCR5* by these autoimmune risk loci.

### SLE-associated rs1432296 acts as a gain-of-function variant to control *REL* expression

Through the analysis of hundreds of non-coding autoimmune risk loci, SCANDAL provides a unique resource to decipher potential downstream molecular mechanisms of non-coding GWAS risk locus by linking them with their *cis*-regulatory target genes. However, CRISPR-SAM cannot resolve allele-specific effects of risk variants within tested CREs. We therefore performed a massively parallel reporter assay (MPRA) in primary B cells to quantify variant-level effects for hundreds of SCANDAL risk variants and determine their consequences on transcriptional activity (**Fig. 6A**). We identified 91 variants with allele-specific effects on transcriptional activity, including 52 that increased and 39 that decreased reporter gene expression (**Fig. 6B**, fig. S9A-B, table S10). Of these variants, 58% (53/91) are predicted to disrupt sequence motifs for transcription factors normally expressed in B cells (fig. S9C, table S11). Notable variant-specific effects included risk variants with experimentally-mapped target genes, such as rs33384 (*NR3C1*), rs12212594 (*HLA-B, HLA-C*), rs6589706 and rs630923 (*CXCR5*), rs4705950 and rs2548998 (*IRF1*), and rs12946510 (*IKZF3*) (**Fig. 6C**, fig. S9D). One of the SCANDAL loci for which we identified variant-specific effects in the lentiMPRA was SLE-associated variant rs1432296 (C/T) which had a gain-of-function effect on reporter activity (**Fig. 6D**) and which is found within a CRE that regulates expression of *REL* (**Fig. 6E-F**). Although MPRA allows large-scale interrogation of variant effects, it lacks endogenous genomic context. We therefore used prime editing to introduce the rs1432296 risk allele (T) in primary human B cells and quantified the relationship between the risk allele frequency and *REL* expression (*24*) (**Fig. 6G**). In fitting with a gain-of-function variant effect for rs1432296, the frequency of T risk allele significantly correlated with increased *REL* expression (**Fig. 6H**). Together, these results provide strong genetic evidence that rs1432296 may contribute to SLE risk by increasing the expression of *REL*.

**Fig. 6.**
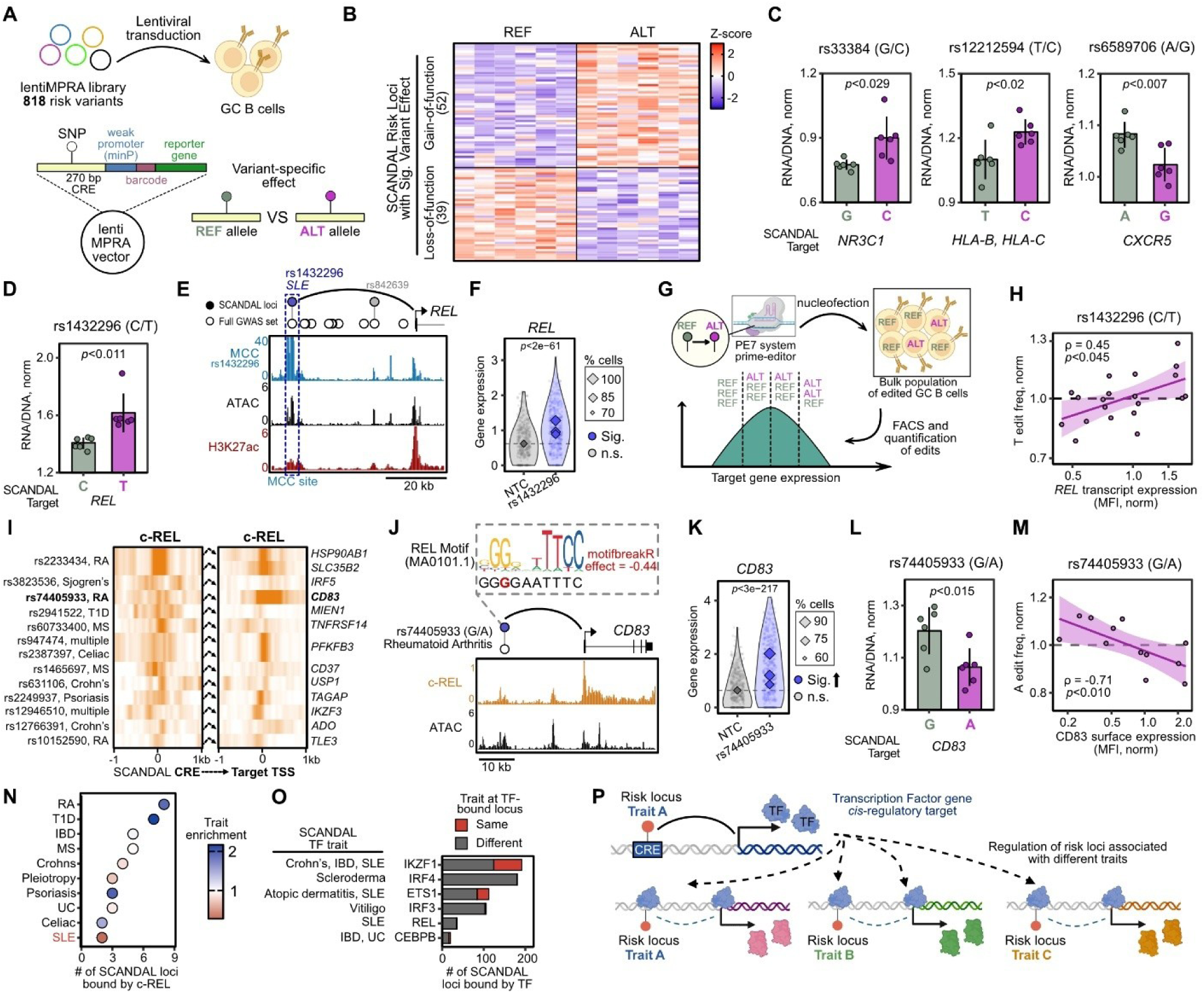
Massively parallel reporter assays and *in situ* prime editing identify loss- and gain-of-function autoimmune risk variant effects at SCANDAL loci. (A) Lentiviral massively parallel reporter assay (lentiMPRA) to quantify variant-specific effects on transcriptional activity. (B) LentiMPRA transcription efficiencies (z-score) for reference (REF) and alternate (ALT) alleles with significant variant-specific effects (*p*<0.05; paired sample t-test; n = 6). (**C**) LentiMPRA transcriptional efficiency (RNA/DNA, norm) for REF and ALT alleles at SCANDAL regions, with *cis*-regulatory target gene annotated below. *p* denotes paired sample t-test (n = 6, ±SD). (**D**) LentiMPRA transcriptional efficiency (RNA/DNA, norm) for REF (C) or ALT (T) alleles at rs1432296. (**E**) Genome snapshot of CRISPR-SAM targeting and Micro-Capture-C (MCC) at rs1432296, including SCANDAL effect on target gene *REL* as curved line. (**F**) *REL* expression after CRISPR-SAM perturbation for cells containing rs1432296-targeting or non-targeting control sgRNAs. Circular points reflect single cell expression values, while diamonds indicate mean expression values for all cells containing individual sgRNAs where size corresponds to percentage of cells in which the gene was detected. *p* denotes Benjamini-Hochberg-adjusted SCEPTRE *p*-value. (**G**) CRISPR-Cas9 Prime-Editing (PE7) in GC B cells to test risk variant-specific effects *in situ* via quantile-based sorting of target expression and amplicon sequencing. (**H**) rs1432296 ALT (T) allele frequency and *REL* transcript mean fluorescence intensity (MFI) in prime-edited GC B cells (n=5). Spearman’s rank correlation (ρ) and *p*-value are shown. (**I**) REL ChIP-seq signal at SCANDAL risk loci and their SCANDAL target promoters. CREs are centred on open chromatin peak or TSS. (**J**) Genome snapshot of REL-bound rs74405933 SCANDAL risk locus, with motifbreakR prediction for A allele effect on REL motif. SCANDAL effect on target gene *REL* as curved line. (**K**) *CD83* expression after CRISPR-SAM perturbation for cells containing rs74405933-targeting or non-targeting control sgRNAs, with formatting as detailed in (F). (**L**) LentiMPRA transcriptional efficiency (RNA/DNA, norm) for REF (G) or ALT (A) alleles at rs74405933. (**M**) rs1432296 ALT (A) allele frequency and CD83 surface expression (MFI) in prime-edited GC B cells (n=5). Spearman’s rank correlation (ρ) and *p*-value are shown. (**N**) Number of SCANDAL loci bound by REL, denoting enrichment of REL-bound loci within each trait as compared to all SCANDAL loci with gene targets for this trait. (**O**) Binding of SCANDAL target transcription factors (TF) to other SCANDAL loci with targets, showing if the bound locus has the same trait as the locus controlling the TF expression. (**P**) Proposed model of genetic pleiotropy underlying pan-disease autoimmune risk. While *cis*-regulatory effects on TF expression may be mediated through a GWAS risk variant associated with Trait A, the downstream targets of that TF include many non-coding risk loci (or their regulatory target genes) with genetic associations for different traits.

### Widespread transcription factor binding mediates pleiotropy across autoimmune risk loci

*REL* encodes the transcription factor c-REL which is a member of the NF-κB family with key roles in B cell survival and maintenance of the GC reaction (*49–51*) and consistent with this CRISPR-SAM activation of the rs1432296 locus increased proliferation of GC B cells (fig. S10A). To explore if c-REL might regulate other autoimmune risk loci, we next analysed c-REL chromatin occupancy data (*48*). We found that c-REL bound to 36 SCANDAL risk loci with *cis*-regulatory targets, as well as promoters of 142 SCANDAL target genes (fig. S10B, table S12). Notably, for 11 risk loci, c-REL occupied both distal CRE and target gene promoter, including for immune-relevant genes such as *TNFRSF14*, *IKZF3* and *CD83* that span diverse autoimmune traits (**Fig. 6I**). We further examined a SCANDAL CRE that regulated expression of *CD83* (a gene important for B-cell activation and maturation) and contains the rheumatoid arthritis-associated variant rs74405933 for which the risk A allele is predicted to disrupt a REL binding motif (**Fig. 6J-K**). Indeed, we found that the risk A allele resulted in decreases in both reporter transcriptional activity (**Fig. 6L**) and CD83 surface protein expression after prime editing was used to introduce the variant *in situ* in primary B cells (**Fig. 6M**). Although rs1432296 (*REL*/c-REL) is associated with increased risk of SLE, we found that REL-bound loci, such as rs74405933 near *CD83,* were associated with a range of autoimmune traits, and not SLE specifically (**Fig. 6N**). To explore how common this type of pleiotropic transcription factor network might be, we first identified transcription factor genes that were SCANDAL *cis*-regulatory targets and expressed in B cells (41 total, fig. S10C). We then examined chromatin occupancy of 6 transcription factors (IKZF1, IRF4, ETS1, IRF3, REL, CEBPB) with known roles in immune cell function. These transcription factors frequently bound distal autoimmune risk loci and/or SCANDAL *cis*-regulatory target promoters (fig. S10D-F) and, as with c-REL, these risk loci bound by transcription factors binding were associated with diverse autoimmune traits that were typically different to the trait at the locus encoding the transcription factor gene (**Fig. 6O**). Together, these observations support a model in which dysregulation of transcription factor genes by non-coding autoimmune risk variants can exert downstream pleiotropic effects at other non-coding risk loci (and their *cis*-regulatory targets) beyond the primary disease association (**Fig. 6P**), and highlight how dysregulated non-coding gene regulatory networks may contribute to shared genetic mechanisms in autoimmunity.

## Discussion

In this study, we experimentally mapped *cis*-regulatory gene targets for hundreds of non-coding risk loci from over 30 autoimmune GWAS traits. By combining gene expression, surface protein expression and CRE transcriptional readouts with a single-cell perturbation screen in primary human B cells, SCANDAL provides a powerful resource to interpret gene regulatory landscapes at autoimmune-associated risk loci. Quantification of SCANDAL variant effects discovered that SLE-associated rs1432296 acts as a gain-of-function variant in primary B cells to increase expression of the NF-ΚB family member *REL*/c-REL to potentially mediate pleiotropy across diverse diseases. Our study therefore marks significant progress in systematically interrogating the consequences of non-coding risk variants in autoimmune disease.

GWAS have identified thousands of variants linked to human traits and disease risk, yet most of them lie in non-coding regions and translating these associations into biological mechanisms has remained a major challenge due in part to difficulty in high-throughput perturbations in primary cells. We overcame limitations in traditional viral transduction protocols to perform high-throughput CRISPR-SAM activation screens in primary human B cells, a cell type that is highly relevant to diverse autoimmune diseases due to the prevalence of self-reactive B cells, autoantibody production or altered B cell-derived cytokine production (*29, 52*). Our SCANDAL resource identified 524 *cis*-regulatory gene target genes regulated by 378 autoimmune risk loci. While the majority of identified CRE-gene pairs were concordant with eQTL-gene pairs (>70%), recent studies have highlighted differences between CRISPRi-based screens and eQTL studies

(*53*). Whether this is related to analytical limitations or systematic differences in evolutionary constraint and regulatory complexity of GWAS risk variants compared to eQTL variants (*54*) is not clear. Chromosome conformation capture assays such as Hi-C, 4C, and Capture-C are widely used to map CRE-target relationships, but the signal from these assays is highly dependent on distance and sometimes making it challenging to identify very distal (where the signal decays to close to background) or very proximal (where physical proximity leads to artificially high interaction frequencies that masks functionally relevant CREs). Our combined analysis of autoimmune risk loci with MCC, which can achieve base-pair resolution (*35*), and CRISPR-SAM perturbation provides a high-resolution, sensitive, flexible and robust approach to map physical and functional variant-to-gene interactions.

Pooled CRISPRi-based perturbation screens rely on being able to measure reductions in transcript abundance and therefore require *cis-*regulatory target genes to be expressed at moderate to high levels. In contrast, we demonstrate that CRISPRa-based screens can map regulatory effects even when targets are weakly expressed or transcriptionally silent. This is particularly important for pooled CRISPR-perturbation screens coupled with single-cell transcriptomic readouts where the sensitivity to detect lowly expressed genes is a major limitation, especially given that important gene classes such as cytokines and transcription factors are often hard to measure in single-cell assays. In total we mapped gene targets for 49.5% of CRISPR-SAM targeted loci compared to 25% or fewer reported with CRISPRi methods (*16, 17, 22, 55*), which we attribute to the ability to detect positive effects of lowly expressed genes and larger perturbation effect sizes (greater signal-to-noise) that we obtained through dual targeting of both dCas9-VP64 and the p65-HSF1 to non-coding CREs. One unexpected result was that SCANDAL enabled identification of CRE-target relationships for genes not normally detected in B cells but that are expressed in disease-relevant tissue, such as *CTRB1/2* in pancreas (with variants associated with type 1 diabetes). These likely reflect poised CRE-promoter relationships and suggests that CRISPR activation perturbations in tractable cellular models enables identification of locus-gene pairs relevant to experimentally challenging cell types or tissue contexts.

In line with previous studies, we found that functional CRE-target gene pairs can act over long genomic distances, with many targets located >100 kb from the associated risk locus. While the nearest gene was often affected, it was frequently not the sole target, demonstrating that linear proximity alone is often inadequate to predict the full set of regulatory targets for GWAS risk loci. These distal CREs typically exist in regulatory landscapes comprising multiple CREs that coordinate to regulate their target gene expression with widespread redundancy such that deletion (CRISPR knockout) or inhibition (CRISPRi) of individual CREs only results in minor or non-detectable effects on target gene expression (*56–59*). In contrast, CRISPRa-based screens of CRE activity appear to be able to overcome this limitation by ectopically increasing CRE activity and reveal their regulatory potential. Additionally, our study reveals that the transcriptional activity of CREs can provide a readout for CRISPR-based perturbation screens to investigate *cis*-regulatory landscapes. SCANDAL enabled us to quantify how increased transcriptional activity at CREs drives transcriptional activity at target gene promoters but also other CREs within the same regulatory landscape. Whether this co-activation is directly mediated through CRE-CRE physical contacts, or indirectly through CRE-promoter-CRE interaction hubs remains to be determined. We anticipate that broader application of perturbation-based screens of CREs will lead to new discoveries about the specificity and cooperativity of gene regulatory mechanisms in controlling normal gene expression and how non-coding genetic variants may disrupt normal gene regulatory processes in disease.

Although the existence of shared genetic risk between different autoimmune diseases has been recognised, understanding how and if genetic risk loci and variants mediate this risk to different diseases and drive shared versus disease-specific mechanisms has remained a persistent challenge. Our findings support several models of shared and pleiotropic genetic effects in autoimmune disease that may also be relevant to other complex genetic diseases. One quarter of autoimmune *cis*-regulatory target genes were controlled by multiple non-coding risk loci. These loci were often associated with different disease traits, indicating convergence of distinct genetic signals on common target genes and pathways. While in some instances, this may relate to inability to resolve variants in high linkage disequilibrium from the statistical fine mapping resources that we leveraged (*25, 26*), it seems likely that independent risk variants exist within the same locus that control the same target, especially for gene targets that we linked with multiple traits such as *HLA-B*, *CD40*, *IL10*, *CXCR5*, and *PTGER4*. Fittingly, we observe that risk loci for different disease traits converge on shared biological processes such as lymphocyte activation, survival, and antigen presentation that provides mechanistic insight into the shared genetic basis between autoimmune diseases and that may help explain overlapping clinical and immunological features. Another potential mechanism for genetic pleiotropy of autoimmune disease is provided by autoimmune risk variants such as rs1432296 at the *REL* locus. By disrupting the expression of transcription factors that then bind dozens or even hundreds of downstream risk loci or target genes, as reported for *NFE2* and *GFIB* in relation to diverse blood cell traits (*17*), these autoimmune risk variants could lead to diverse changes to cellular identity or function relevant to loss of tolerance in autoimmunity. Finally, our investigation of CRE transcriptional activity at autoimmune risk loci proposes a new model for genetic pleiotropy whereby allele- and disease-specific effects at one CRE could alter the activity of nearby CREs linked with genetic risk for a different autoimmune disease. At loci such as *CXCR5*, this type of non-coding regulatory landscape could result in shared molecular outcomes driven by independent variants, although significant further work is needed to test this model experimentally.

Taken together, our work establishes a vital resource for autoimmune disease genetics and provides conceptual advances into complex genetic diseases more broadly by offering mechanistic insights into function of hundreds of GWAS loci. Our study also provides an important and necessary advance in the transition of large-scale single-cell CRISPR screening from cancer cell lines to more relevant primary cell types, which may prove vital to transition from target gene discovery to translational or clinical pathways.

## Methods

### Ethical approval for use of human biospecimens

Paediatric tonsil samples were collected from children undergoing routine tonsillectomy at Royal Children’s Hospital (RCH), Melbourne, Australia with informed written consent and protocols approved by RCH HREC (88144/RCHM-2022) and processed at the Walter and Eliza Hall Institute (WEHI) under WEHI HREC approval 23/49. Peripheral blood samples were obtained with ethical approval by venesection from volunteers recruited through either the Volunteer Biospecimen Donor Registry (WEHI HREC Project 24/48, WEHI HREC Project 10/02) or the Investigation of Haematopoiesis in Healthy Volunteers Study (CUREC1 R88300/RE001).

### Tonsillar human germinal centre (GC) B cell isolation and culture

Whole paediatric tonsils were collected in RPMI 1640 medium supplemented with 5% heat-inactivated FBS and processed immediately. Tonsillar tissue was dissociated using the gentleMACS™ Dissociator, and mononuclear lymphocytes isolated by Ficoll-Paque™ density gradient centrifugation. After cell counting and determination of viability with TrypanBlue staining, tonsillar mononuclear cells were cryopreserved in 90% FBS and 10% DMSO until further use. GC B cells were isolated via negative selection with human B Cell Isolation Kit II (Miltenyi Biotec 130-091-151) with the addition of biotin anti-human IgD (IA6-2; BioLegend) and biotin anti-human CD32 (FUN-2; BioLegend) antibodies at final amounts of 250 ng and 62.5 ng per 100 M total tonsillar mononuclear cells. Isolation purity was assessed by flow cytometry with anti-human IgD (IA6-2; BioLegend), CD27 (O323; BioLegend), CD32 (FUN-2; BioLegend), CD38 (HB-7; BioLegend), CD138 (DL-101; BioLegend) and CD19 (HIB19; BioLegend) antibodies and analysed on the Cytek Aurora flow cytometer. Purified tonsillar GC B cells were plated on a layer of irradiated YK6 feeder cells expressing CD40Lg and IL21 (YK6-CD40Lg-IL21) at a ratio of 6.7:1 and cultured in RPMI 1640 medium supplemented with 20% heat-inactivated FBS, 2 mM GlutaMAX, 25 mM HEPES, 1 mM sodium pyruvate, 100 µg/mL Normocin, and 1% Pen-Strep as described (*60*).

### Total B cell isolation from peripheral blood

Isolated veinous blood (100 mL) was mixed in a 1:1 ratio with room temperature phosphate buffered saline (PBS) before layering 30 mL on 15 mL Lymphoprep™ (STEMCELL Technologies). Peripheral blood mononuclear cells were isolated by spinning this mixture for 30 min at 800 × *g* with minimal break and acceleration, before enrichment of CD19^+^ B cells by positive selection with CD19 MicroBeads (Miltenyi Biotec) following the manufacturer’s protocol.

### Cloning of CRISPR-SAM components

The blasticidin resistance gene in LentiSAMv2 (a gift from Feng Zhang; Addgene #75112 (*61*)) was replaced with Cherry Fluorescent Protein (ChFP) by digesting LentiSAMv2 with BsrGI-HF (New England Biolabs, NEB) and EcoRI-HF (NEB) and replaced with the ChFP gene amplified from pLVX-EF1α-IRES-mCherry vector (Takara Bio 631987) to generate *LentiSAMv2-ChFP*. The subsequent dCas9-VP64-T2A-ChFP fragment was amplified and cloned into retroviral vector MSCV-BCL6-t2A-BCL2 (Addgene #135305; (*60, 62*)) digested with XhoI (NEB) and SalI-HF (NEB) to replace the BCL6-T2A-BCL2-IRES-CD2 fragment and generate *MSCV-dCas9-VP64-ChFP*. Similarly, MCP-p65-HSF1-T2A-GFP was amplified from the MS2-P65-HSF1_GFP vector (a gift from Feng Zhang; Addgene #61423 (*31*)) and cloned into MSCV-BCL6-t2A-BCL2 backbone to generate *MSCV-MCP-p65-HSF1-GFP*. ChFP in the lentiviral CROP-sgRNA-MS2 vector (a gift from Wolf Reik; Addgene #153457) was replaced with BFP by GenScript using their gene synthesis service to generate *CROP-sgRNA-MS2-BFP*. PCR fragments for in-house cloning were amplified using NEBNext Ultra II Q5 Master Mix (NEB) and ligations performed with NEBuilder® HiFi DNA Assembly Master Mix (NEB). Individual single-guide RNAs (sgRNAs) for CRISPR activation were designed with CRISPick (Broad Institute) (table S13A). sgRNA sequences were cloned between BsmBI restriction sites in CROP-sgRNA-MS2-BFP (GenScript).

### Preparation of RD114-pseudotyped retrovirus and GALV-pseudotyped lentivirus

For retrovirus preparation, HEK-293T cells were plated at a density 12.5×10^6^ cells in T150 flasks 16 h before transfection with Lipofectamine 3000 (Thermo Fisher Scientific) according to manufacturer’s instructions. Total 47.4 µg of DNA containing a molar ratio of 1:1:12 of the RD114 envelope plasmid (a gift from Daniel Hodson), gag-pol packaging plasmid (a gift from Marnie Blewitt) and transfer plasmid (MSCV-dCas9-VP64-ChFP, MSCV-MCP-p65-HSF1-GFP, or MSCV-BCL6-t2A-BCL2) was used for transfection. Supernatants at both 24 and 52 h post-transfection were collected and pooled. MSCV-dCas9-VP64-ChFP viral supernatants were further concentrated 30x times with Lenti-X concentrator (Scientifix) according to manufacturer’s instructions. Viral supernatants were stored at −80°C until required. Lentivirus was prepared similarly to retrovirus, except that 19×10^6^ HEK-293T cells were plated per T150 flask and a 1:3:4 molar ratio of GaLV envelope plasmid (a gift from Daniel Hodson; Addgene #163612), psPAX2 packaging plasmid (a gift from Didier Trono; Addgene #12260), and transfer plasmid (CROP-CD19-MS2-BFP, CROP-CD86-MS2-BFP, CROP-CD5-MS2-BFP, CROP-CD83distal-MS2-BFP, CROP-ntc-MS2-BFP, CROP-SCANDAL-MS2-BFP) (total 47.4 µg) for transfection.

### Viral transduction for CRISPR-SAM activation in human GC B cells

Purified tonsillar GC B cells were plated on a layer of irradiated YK6-CD40Lg-IL21 feeder cells and cultured for two days. For CRISPR-SAM activation, the media was then replaced with 1.5 mL of pooled viral supernatant comprised of 0.7 mL of CROP-sgRNA-BFP lentivirus, 0.7 mL of MSCV-MCP-p65-HSF1-GFP retrovirus and 0.1 mL of 30x concentrated MSCV-dCas9-VP64-ChFP retrovirus followed by spinfection at 1,500 × g for 3 h at 32°C. Polybrene (Merck) and HEPES (Thermo Fisher Scientific) were added to a final concentration of 10 µg/mL and 25 mM, respectively. After spinfection, the viral supernatant was changed for 1 mL of RPMI 1640 medium supplemented with 20% heat-inactivated FBS, 2 mM GlutaMAX, 25 mM HEPES, 1 mM sodium pyruvate, 100 ug/mL Normocin and 1% Pen-Strep. Cells were grown for 3 days and collected by manual scraping before filtering through a Falcon 70 µm Cell Strainer. Cells were incubated with FcR Blocking Reagent (Miltenyi Biotec) and stained for 20 min in the dark at 4°C with either CD19-APC (HIB19, BioLegend), CD86-APC (IT2.2, BioLegend), CD5-APC (UCHT2, BioLegend), or CD83-APC (HB15e, BioLegend), prior to analysis on the Cytek Aurora spectral flow cytometer to quantify ChFP^+^ (dCas9-VP64), GFP^+^ (MCP-p65-HSF1), and BFP^+^ (sgRNA) cells. Spectral data were unmixed using SpectroFlo (Cytek) prior to analysis with FlowJo v10.8.1 to quantify APC median fluorescence intensities (MFI) for triple-positive cells (GFP^+^;ChFP^+^;BFP^+^) representing cell populations carrying all 3 CRISPR-SAM components for either target sgRNA or NTC sgRNA controls.

### Curation of fine-mapped GWAS risk variants across diverse autoimmune traits

Full GWAS catalogues with statistically fine-mapped SNPs were downloaded from PICS2 (version 2024-04-19) (*26*) and OpenTargets Genetics (latest v2d dataset accessed on 17.03.2024) (*25*) and all lead SNPs associated with autoimmune disease traits were retained. Risk variants with ambiguous GWAS trait definitions such as “Celiac disease or Rheumatoid arthritis” were re-classified as “Pleiotropy”, and redundant GWAS traits were consolidated into 31 autoimmune disease traits with a total of 5655 unique risk variants (table S1). The trait dictionary containing full original trait definition as well as the SCANDAL assigned trait is available in table S14. SNPs were assessed whether they overlapped with the MHC/HLA super-locus (chr6:28510120-33480577; ncbi.nlm.nih.gov/grc/human/regions/MHC?asm=GRCh38). Risk variants associated with multiple non-redundant traits (excluding traits Granulomatosis, Pernicious anemia, and Polymyalgia rheumatica due to limited multi-trait variants) were used in force-direct graph analysis with a Fruchterman−Reingold Layout to visualise shared risk associations between traits.

### Prioritisation and annotation of risk variants using functional genomic datasets

SNP genomic locations and distances to the nearest transcription start site (TSS) were determined with HOMER v5.1 using annotatePeaks.pl function and hg38 gtf file filtered for protein-coding transcripts (table S1). Population allele frequencies were uploaded from 1000 Genomes Project. To prioritise autoimmune risk variants with regulatory potential in human B cells, single-cell ATAC peaks from naive B cells, activated B cells, GC B cells, memory B cells and plasma cells were obtained (*14*). Bulk GC B cells ATAC-seq datasets (EGAD00001002917; *63*) were aligned to GRCh38 with bowtie2 v2.4.4 (*64*) before retaining uniquely mapped reads and duplicate read removal with samtools v1.16.1 (*65*), peakcalling with MACS2 (*66*) and replicate peaks with p-values <0.05 and FE >3.5 were merged using bedtools v2.31.1 (*67*). The resulting 684 open chromatin regions containing autoimmune SNPs were designated CREs (*cis*-regulatory elements), and 818 risk variants overlapping with 684 B cell CREs were prioritised for further testing (table S2).

Biological replicate bam files were downsampled to have equal numbers of reads and merged with samtools before generating representative genome tracks with deepTools v3.5.1 (*68*) and visualised with the UCSC Genome Browser (*69, 70*). Bulk H3K27ac, H3K4me1, H3K4me3, and H3K27me3 ChIP-seq datasets from GC B cells (EGAD00001002442; *63*) were processed similarly. Read counts were obtained using multicov from bedtools v2.31.1 (*67*) and normalised to total number of mapped reads to calculate counts per million (CPM). CRE genomic locations were determined with HOMER v5.1 using annotatePeaks.pl and hg38 gtf file filtered for protein-coding transcripts. Promoter CREs were defined as open chromatin regions that overlapped with annotated transcription start sites based on HOMER annotation, or that were enriched for the promoter-associated H3K4me3 relative to enhancer-associated H3K4me1 ((H3K4me3 CPM - H3K4me1 CPM) > 3). All other open chromatin regions were considered as distal CREs (table S2). Heatmaps of chromatin accessibility and histone modification signals centred on open chromatin peaks were produced using deepTools v3.5.1 (*68*). Fisher Exact tests were used to determine trait enrichment and depletion for risk variants before and after this functional prioritisation.

### CRISPR-SAM sgRNA library design for screens of non-coding risk loci

Autoimmune risk loci for CRISPR-SAM activation were defined as ±100 bp regions centred on statistically fine-mapped autoimmune risk variants that overlapped with B cell open chromatin. In a limited number of cases where two variants were located within 100 bp of each another, the 200 bp window was centred equidistant between both variants. sgRNA sequences for each 200 bp locus were designed with CRISPick (mechanism = CRISPRa, enzyme = SpyoCas9 NGG (Chen), quota = 20, report unpicked sgRNAs, allow picking guides with MAX off-target matches) (*71, 72*). After removing guides with Pick Order > 6, top 3 sgRNAs with highest On-Target Efficacy Scores were selected for each locus. A limited number of loci (28) were excluded due to poor targetability where only 1 out of 3 sgRNAs had On-Target Efficacy Score > 0.2, and a small number of sgRNAs (4) with low On-Target Efficacy Score (<0.2) and high Off-Target Rank were manually replaced by sgRNAs with higher Off-Target Rank. We considered the spacing of sgRNAs within each 200bp window, and in some cases where the cut sites of all 3 sgRNAs were within 4 bp of each other, replacement sgRNAs with PAM targeting sites located further away within the same window were selected. As controls, we designed sgRNAs for 10 loci with no TSS within 500 kb and no open chromatin signal in B cells within 30 kb (gene desert controls), 20 loci within 5-7 kb of any TSS and no open chromatin signal in B cells within 3 kb (distance-based controls), and three additional loci including *CD5* and *CD19* promoters and a non-coding regulatory element encoding rs74405933 upstream of CD83, in addition to 18 non-targeting sgRNAs from Calabrese Human CRISPR Activation Pooled Library (*72*). In total, our CRISPR-SAM library comprised 2,403 sgRNA sequences targeting 763 autoimmune risk loci, 20 distance-based controls, 10 gene desert controls, 2 TSS controls and 18 non-targeting controls (table S3).

### Cloning of the sgRNA library into CROP-sgRNA-MS2-BFP vector

An oligonucleotide pool of sgRNA sequences flanked by 5’ (AGGCACTTGCTCGTACGACGCGT CTCACACCG) and 3’ (GTTTCGAGACGTTAAGGTGCCGGGCCCACAT) adapters was synthesised by Twist Bioscience, and 10 ng of this oligonucleotide pool was amplified with 0.5 µM sgRNA_lib_FW and sgRNA_lib_RV primers (table S13B) using the NEBNext Ultra II Q5 Master Mix (NEB) and the following cycling: 3 min at 98°C, (98°C for 30 s, 60°C for 30 s, 72°C for 30 s) × 9 cycles, 72°C for 1 min. PCR product was purified with DNA Clean & Concentrator-5 kit (Zymo Research) and cloned into CROP-sgRNA-MS2-BFP vector pre-digested with BsmBI-v2 (NEB) via Golden Gate Assembly with 150 ng pre-digested vector, 5 ng purified PCR product (5 ng), 1X T4 DNA Ligase Reaction Buffer (NEB), NEBridge® Golden Gate Assembly Kit (BsmBI-v2) (NEB) and following cycling conditions: (42°C for 5 min, 16°C for 5 min) × 60 cycles, 60°C for 5 min. The ligation product was purified with DNA Clean & Concentrator-5 kit and transformed into Endura Electrocompetent Cells (Lucigen). Transformed cells were grown overnight with 100 µg/mL ampicillin (Sigma-Aldrich) and pooled library DNA was purified with PureYield Plasmid Maxiprep System (Promega) prior to sequencing with Oxford Nanopore PromethION 2 Solo to ensure sgRNA library representation.

### Bulk CRISPR-SAM screen for B cell proliferation and survival

Human GC B cells were purified by negative selection from cryopreserved tonsillar mononuclear cells from two donors and cultured as described above. To generate cells stably expressing CRISPR-SAM system, GC B cells were transduced with RD114-pseudotyped MSCV-BCL6-t2A-BCL2 retrovirus (day 2) and subsequently transduced with RD114-pseudotyped MSCV-dCas9-VP64-ChFP and RD114-pseudotyped MSCV-MCP-p65-HSF1-GFP retroviruses (day 8) followed by sorting for ChFP^+^GFP^+^ cells (day 15). CRISPR-SAM library was then delivered one week later (day 22) via GaLV-pseudotyped CROP-sgRNA-BFP lentivirus, and a baseline sample was collected 3 days later (day 25) before continued culture and expansion of all remaining cells for a further 12 days (day 37). gDNA extractions were performed with Quick-DNA Midiprep Plus Kit (Zymo Research), with the entirety of each gDNA sample used to amplify sgRNAs in multiple 50 µL reactions containing 5 µg gDNA each using the NEBNext Ultra II Q5 Master Mix (NEB), 0.5 µM P5_mix_FW and P7-CROPa_A0X_RV (table S13B) and following conditions: 95°C for 3 min, (95°C for 15 s, 60°C for 30 s, 72°C for 30 s) × 28 cycles, 72°C for 7 min. PCR products were purified via 1X AMPure XP bead cleanup, sequenced with the Ligation Sequencing Kit V14 (SQK-LSK114) on the Oxford Nanopore PromethION 2 Solo platform. Basecalling was performed with super-accurate model (dna_r10.4.1_e8.2_400bps_sup@v5.0.0) using dorado (v0.7.3), reads were trimmed and quality-filtered (q>20) using cutadapt (*73*). To identify sgRNA sequences that became enriched in the B cell over the 12 day expansion period, sgRNA count tables were made and differential testing was performed with MAGeCK (*74*) using default settings.

### High-throughput CRISPR-SAM screen in human GC B cells with single-cell readout

Primary human tonsillar GC B cells were isolated by negative selection from cryopreserved tonsil mononuclear cells from one male (6 y.o.) and one female (3 y.o.) donor and cultured on irradiated YK6-CD40Lg-IL21 feeder cells as described above. Pooled single-cell CRISPR-SAM transduction and analysis was performed on these two biological samples (6M, 3F) on two independent occasions (R1, R2), as follows: On day 2 post-isolation cells were transduced with RD114-pseudotyped MSCV-BCL6-t2A-BCL2 to limit plasma cell differentiation, followed by tripartite transduction with GaLV-pseudotyped CROP-sgRNA-BFP library, RD114-pseudotyped MSCV-MCP-p65-HSF1-GFP and RD114-pseudotyped MSCV-dCas9-VP64-ChFP of 2.9-10.9 × 10^6^ cells on day 8 as described above. Cells were harvested three days post-transduction (day 11 since isolation) and incubated with Human TruStain FcX Fc Receptor Blocking Solution (BioLegend) before being split into 5 tubes per donor. Each tube was stained with the TotalSeq-C Human Universal Cocktail, V1.0 (BioLegend) (4-6 µL per 1 M cells) and a unique TotalSeq-C hashtag oligonucleotide (HTO) per tube, prior to pooling and live-dead staining with propidium iodide (Miltenyi Biotec). Triple-positive CRISPR-SAM (GFP^+^;ChFP^+^;BFP^+^) cells were sorted on the BD FACS Aria into FACS buffer (PBS, 2% heat-inactivated FBS) and immediately processed for single-cell capture with the Chromium GEM-X Single Cell 5’ Reagent Kit v3 (10X Genomics). Cells were superloaded at 54,000-66,400 cells per lane (5 lanes R1; 2 lanes R2), followed by gene expression (GEX), antibody-derived tags (ADT), hashtag oligonucleotides (HTO) and CRISPR Guide Capture (only for R2) library construction according to manufacturer’s protocols. For sgRNA detection, we leveraged CROP-seq vector properties (*75*) in which sgRNAs are transcribed from both U6 and EF-1α promoters allowing them to be amplified from polyadenylated cDNA libraries. For each library, 200 ng 10X GEX cDNA was amplified with NEBNext Ultra II Q5 Master Mix (NEB) and 0.5 µM gRNA_10X_Capt_FW and gRNA_10X_Capt_RV primers (table S13B) (3 days post-transduction (day 11 since isolation)) across 8 × 50 µL reactions with the following conditions: 98°C for 45 s, (98°C for 10 s, 66°C for 75 s) × 19 cycles, 72°C for 4 min. The resulting 2 kb PCR products (containing 10X cell barcode, UMI and sgRNA spacer sequence) were purified with 0.6X AMPure XP beads (Beckman Coulter) and 500 ng used to prepare Oxford Nanopore libraries with Native Barcoding Kit 24 v14 (Oxford Nanopore SQK-NBD114.24). The resulting sgRNA libraries were sequenced with Oxford Nanopore PromethION 2 Solo, while all other libraries were sequenced on the Illumina NovaSeq X Plus with 150bp paired-end reads and the following depths: ∼37,000 GEX reads per cell, ∼5,000 sgRNA reads per cell (10X CRISPR Capture; R2 only), ∼9,800 ADT+HTO reads per cell.

### Single-cell CRISPR-SAM data processing and quality control

For Illumina sequencing, bcl files were converted into fastq using bcl2fastq v2.20.0 (Illumina), while Oxford Nanopore pod5 files underwent basecalling with the super-accurate model (dna_r10.4.1_e8.2_400bps_sup@v5.0.0) of dorado v0.7.3 (Oxford Nanopore Technologies) and resulting fastq files were trimmed and quality-filtered (q>20) using cutadapt (*73*). Cellranger multi (v8.0.1; 10X Genomics) was used to align GEX libraries to the hg38 and generate UMI count matrices for GEX, ADT, HTO and 10X sgRNA libraries, while long-read sgRNA sequencing libraries were processed with custom scripts to generate cell barcode-sgRNA UMI count matrices (all sgRNAs with UMI ≥ 3 were assigned to a cell barcode). Cell barcode-count matrices were then analysed with Seurat (v5.1.0) (*76*) in R (v4.4.1). Cell barcodes with high mitochondrial gene expression (>7.5%) were removed, prior to log10 normalisation of GEX count matrices and centred log ratio (CLR) normalisation for HTO and ADT counts. HTOs were demultiplexed using HTODemux function (assay = “HTO”, positive.quantile = 0.99, kfunc=’clara’) to identify biological donor samples and remove technical doublets from superloading the 10X capture, before integration of all captures were integrated using reciprocal principle component analysis (RPCA; IntegrateLayers). After quality control and data integration, we retained data for 256,790 cells, with 201,191 cells having an assigned sgRNA, with a median of 116 cells per sgRNA and on average 1.4 unique sgRNAs detected per cell.

### Identification of cis-regulatory targets of CRISPR-SAM perturbations with SCEPTRE

We used SCEPTRE (v0.10.0; *33*) to test for differentially expressed genes and cell surface proteins in response to CRISPR-SAM activation of autoimmune risk loci. GEX and ADT count matrixes were generated from the integrated final Seurat object with DropletUtils (v1.26.0; *77, 78*)). SCEPTRE differential gene expression analysis was performed for all genes (GEX) (table S5) or cell surface proteins encoded by genes (ADT) (table S6) within ±500 kb of each targeted locus with high MOI settings, combining cells that contained any of the 3 sgRNAs per target locus and using cells with non-targeting sgRNAs as the control group. We found no major difference in the number of *cis*-regulatory targets identified for high- or low-MOI settings (data not shown). Following this pooled analysis, we quantified the effects from individual sgRNAs for each locus by repeating SCEPTRE analysis with a singleton sgRNA strategy. We then retained only results that were significant and concordant for ≥2 sgRNAs (SCEPTRE *p* < 0.05) (table S4).

### Quantification of *cis*-regulatory element transcription in single-cell CRISPR-SAM screens

Cellranger BAM files from 10X 5’ scRNA-seq GEX libraries were used for the single-cell analysis of five-prime end (SCAFE) v1.0.0 (*45, 79*) to identify and quantify 5’ TSS signal at CREs. Briefly, scafe.tool.sc.bam_to_ctss was used with default settings to identify 5’ non-encoded guanines (reflecting the 5’ mRNA cap) and extract the precise 5’ position of GEX reads, while considering strand invasion artifacts through alignment of template-switch oligo sequence to immediate upstream sequences. Identified TSS were then merged into tCREs using scafe.tool.cm.annotate with default parameters, and tCRE UMI/cell-barcode count matrixes were generated using scafe.tool.sc.count. 5’TSS signals were then quantified as a sum of UMI counts within 500 bp regions around each SNP in cells containing either NTC sgRNAs (basal level) or targeting sgRNAs (after activation) (table S7), and scafe.tool.cm.ctss_to_bigwig was used to generate bigWig tracks.

To quantify total RNA signal at distal CREs, we processed bam files (generated by cellranger) with dropEst (*80*) using the default parameters and a custom gtf file with coordinates of all B cell open chromatin peaks within 500 kb of each tested locus. The resulting matrixes were used for differential CRE expression analysis with SCEPTRE (v0.10.0; *33*). The testing was done for all open chromatin regions (response CREs) within 100 kb of activated locus with high MOI settings, combining cells that contained any of the 3 sgRNAs per target locus and using cells with non-targeting sgRNAs as the control group. We then retained results that were called as significant by SCEPTRE (p.adj < 0.1). To identify distal CREs with significant increase in transcription upon CRISPR activation, for each tested perturbation we selected the response CRE containing the sgRNA-targeted locus (table S8). To identify intergenic CRE-CRE coregulation pairs (table S9), we masked CREs overlapping protein-coding transcripts (GRCh38.p14) with bedtools v2.31.1 (*67*).

### Cell type- and disease-specific patterns of target gene expression

Single-cell gene expression data for 13 different immune cell types was taken from a published study (*14*). For a given gene, a percentage of cells with non-zero counts was quantified for each cell type and visualised on a heatmap. A gene was considered to B cell enriched if the percentage of non-zero cells was >60% in total B cell population (Naïve B, Activated B, GC B, Memory B, Plasma B) and <30% in total non-B cell population. Alternatively, genes with <30% non-zero cells in total B cell population and >60% non-zero cells in total non-B cell population were assigned as “non-B enriched”. Finally, genes with <10% non-zero cells for each of 13 immune cell types were classified as “non-detected”, while all remaining genes were considered “widely expressed”. Differential expression values between autoimmune disease cohorts (cases/controls) for the 378 SCANDAL *cis*-regulatory target genes were downloaded from the Autoimmune Disease Explorer (*81*). Specifically, peripheral blood-derived expression values were compared with disease-relevant tissues such as salivary gland for Sjogren’s syndrome, skin for systemic sclerosis, and synovial joint for rheumatoid arthritis.

### Comparative analysis with CRISPRi-based studies and eQTL resources

Gene Onthology analysis was performed with Metascape (*82*). eQTL data for 11 different tissues was downloaded from EMBL-EBI eQTL catalogue (QTD000356, QTD000326, QTD000266, QTD000271, QTD000296, QTD000171, QTD000281, QTD000291, QTD000311, QTD000331, QTD000346) (*83*). All SNP-gene pairs with *p*-value < 0.05 were considered eQTLs. For the effect sizes comparison between CRISPRi and CRISPRa screens, we downloaded analysed data from published studies (*16, 17, 20–22, 55*). Threshold values for statistical significance were used as described in original studies; otherwise, the following values were used: Alda-Catalinas et al. (table S2 from original study, FDR < 0.05), Green et al. (table S3A, Hit == TRUE), Ho et al. (tabl S21-22, significant == TRUE), Gasperini et al. (table S1), Morris et al. (table S3F, Skew.fit.p.value <= 0.0001), Chardon et al. (table S10, Benjamini-Hochberg adjusted *p*-value < 0.1).

### Micro Capture-C Library Preparation and Sequencing

For all Micro Capture-C (MCC) experiments, peripheral blood-derived B cells were cultured for 3 days at a starting density of 2×10^5^ cells per well of a 24 well plate using the ImmunoCult Human B cell Expansion kit (STEMCELL Technologies) according to the manufacturer’s instructions prior to MCC library generation which was performed as previously described (*35, 84*). In brief, 1×10^7^ B cells were fixed at room temperature for 10 min with 2% final concentration of formaldehyde. The reaction was quenched with ice-cold 1M glycine and spun at 400 × *g* for 5 min. Cells were resuspended in 1 mL of PBS before permeabilization with 0.005% digitonin (Sigma-Aldrich) for 15 min at room temperature. Cells were snap frozen on dry ice and stored at −80°C.

Frozen cell aliquots were thawed to room temperature before being split into three separate digestion reactions with different amounts of MNase (typically 5-30 Kunitz U) in a reduced-calcium MNase buffer (10 mM Tris-HCl (pH 7.5), 1 mM CaCl_2_). Reactions were incubated for 1 h at 37°C with constant mixing (550 rpm) prior to quenching with ethylene glycol-bis(2-aminoethylether)-N,N,N′,N′-tetraacetic acid (EGTA) to a final concentration of 5 mM and cell pelleting and resuspension in PBS containing 1% EGTA. 10 % of each reaction was stored as a digest control. Remaining cells were pelleted and then resuspended in DNA ligase buffer (NEB) supplemented with 400 µM dNTPs and 5 mM EDTA before addition of DNA Polymerase I large (Klenow) fragment (NEB) to 100 U/µL, T4 polynucleotide kinase (NEB) to 200 U/µL and T4 DNA ligase (Thermo Fisher Scientific) to 300 U/µL. This reaction was incubated for 2 h at 37°C, then for 8 h at 20°C with constant mixing at 550 rpm, then held at 4°C overnight. Ligated chromatin as well as digest controls had crosslinks reversed at 65°C in the presence of proteinase K (Qiagen), before DNA purification with the DNeasy Blood and Tissue Kit (Qiagen) followed by quality control of digestion and ligation with the D1000 ScreenTape Assay (Agilent).

Selected MCC libraries were sonicated to 200 bp average fragment size before library preparation with the NEBNext Ultra™ II DNA Library Prep Kit for Illumina (NEB). After indexing MCC libraries were pooled together, lyophilised using a vacuum centrifuge at 55°C and resuspended in hybridization buffer according to the HyperCapture Target Enrichment Kit (Roche) protocol. Biotinylated oligonucleotides were designed with Capsequm (github.com/jbkerry/capsequm) and synthesised as a pool of xGen Custom Lockdown Probes (IDT). 22 µL of a 16X pool was added to the capture reaction before incubation at 47°C for 72 h. Reactions were then washed according to the Roche HyperCapture instructions, prior to a second capture reaction for 24 h to increase target recovery. Libraries were sequenced with 150 bp paired-end reads on the Illumina NovaSeq X Plus platform to 500,000 reads per capture viewpoint per library.

### Micro Capture-C analysis

MCC analysis was performed as previously described using the MCC pipeline (github.com/jojdavies/Micro-Capture-C) (*35, 84*). In brief, adapter sequences were removed using TrimGalore (v.0.3.1; github.com/FelixKrueger/TrimGalore) and reads were reconstructed using FLASH (v.1.2.11; *85*) before mapping with the non-stringent aligner BLAT (v.35; *86*) to 800 bp genomic windows centred on the capture oligonucleotides used. Reads were separated into different files based on which oligonucleotide reference sequence they mapped to using custom script MCCsplitter.pl before being alignment to the hg38 genome with Bowtie2 (v.2.3.5; *64*), before custom script MCCanalyser.pl was used to remove PCR duplicates and identify ligation junctions. MCCsplitter.pl and MCCanalyser.pl can be accessed for academic use via the Oxford University Innovation software store (https://process.innovation.ox.ac.uk/software/p/16529a/micro-capture-c-academic/1). Data were merged and converted into bigwigs for visualisation on the UCSC Genome Browser (*69, 70*). Tracks presented in this manuscript are pooled averages of 6 replicates derived from 2 technical replicates for 3 biological donors.

### Massively Parallel Reporter Assay (MPRA) design and cloning

An oligonucleotide pool of 270 bp candidate CREs and control sequences (table S15) flanked by 5’ (AGGACCGGATCAACT) and 3’ (CATTGCGTGAACCGA) adapters was synthesised by Twist Bioscience to perform a massively parallel reporter assay (MPRA) as described previously (*87*). Genomic sequences centred on autoimmune GWAS risk variants overlapping B cell open chromatin (see above for details) for both reference (REF) and alternative (ALT) alleles were synthesised, in addition to 150 negative control sequences that lacked open chromatin in B cells (100 regions located 5-7 kb away from nearest TSS and lacking B cell open chromatin peaks within 3 kb; 50 gene desert regions located >500 kb from nearest TSS and lacking B cell open chromatin peaks within 30 kb), 20 scrambled sequences and 2 positive controls (CMV and EF-1α promoters).

To clone the MPRA library into lentiviral vector pLS-mP (a gift from Nadav Ahituv; Addgene #81225), 125 ng of the oligonucleotide pool DNA was amplified in 4 × 50 µL reactions with 0.5 µM MPRAlib_PCR1_FW and MPRAlib_PCR1_RV primers and following conditions: 98°C for 1 min, (98°C for 10 s, 60°C for 15 s, 72°C for 20 s) × 5 cycles, 72°C for 5 min. 150 ng of purified PCR product split into 4 × 50 µL reactions and amplified with 0.5 µM MPRAlib_PCR2_FW and MPRAlib_BC_PCR2_RV primers under following conditions: 98°C for 2 min, (98°C for 15 s, 60°C for 20 s, 72°C for 30 s) × 4 cycles, 72°C for 5 min. At this step, a unique 13 bp molecular barcode (from primer MPRA_PCR2_FW) was added to each CRE. The purified product from the second PCR reaction (120 ng) was used in a Gibson assembly reaction with 1 µg pLS-mP linearised with Sbfi-HF (NEB) and AgeI-HF (NEB). The purified ligation product was transformed into Endura Electrocompetent Cells (Lucigen). Transformed cells were grown overnight with 100 µg/mL ampicillin (Sigma-Aldrich), and plasmid DNA was purified with PureYield Plasmid Maxiprep System (Promega). This first MPRA library pool was amplified with 0.5 µM MPRAlib_ass_FW and MPRAlib_ass_ RV primers under following conditions: 98°C for 1 min, (98°C for 15 s, 60°C for 20 s, 72°C for 180 s) × 12 cycles, 72°C for 5 min. The resulting amplicon was purified with 1X AMPure XP beads before library construction with the Ligation Sequencing Kit V14 (SQK-LSK114) and Oxford Nanopore sequencing on the PromethION 2 Solo to confirm library representation and quality. To prepare the final library, the first MPRA library pool was digested with PmeI (NEB) and AsiSI (NEB), and 1 µg of purified DNA was used in a Gibson assembly reaction with 30.9 ng of minP dsDNA synthesised as a gBlock Gene Fragment (IDT). The purified ligation product was transformed into Endura Electrocompetent Cells and grown overnight before plasmid DNA isolation, Nanopore library preparation and sequencing as above. Unless otherwise stated, all cloning steps were performed with NEBNext Ultra II Q5 Master Mix (NEB) and purified with DNA Clean & Concentrator-5 kit (Zymo Research) before quantification with Qubit™ 1X dsDNA HS Assay Kit (Invitrogen). Primer sequences are available in table S13B. The final MPRA library was packaged into GaLV-pseudotyped lentivirus as described above and stored at −80°C.

### Lenti-MPRA barcode association

For barcode-CRE association, final MPRA plasmid library was processed and sequenced with Oxford Nanopore sequencing as described above. pod5 files were basecalled using dorado (v0.7.3) and super-accurate model (dna_r10.4.1_e8.2_400bps_sup@v5.0.0), and the resulting fastq files were trimmed and quality-filtered (q>20) using cutadapt (*73*). CRE identification was performed with cutadapt by finding the reads with a perfect match to unique 31 bp CRE sequence (centred on SNP), with 0 error tolerance to ensure distinction between CREs with REF and ALT nucleotides. Barcode, CRE and UMI sequences were extracted for each read, and UMI counts for barcode-CRE pairs were quantified with custom scripts. To assign a barcode to a given CRE, the barcode-CRE pair was required to have ≥3 UMI counts. The uniqueness of each barcode was quantified as number of reads for a given CRE divided by the total number of reads for this barcode. Only barcodes with >95% uniqueness were retained for the final barcode-CRE associations, with a median of 610 barcodes per CRE.

### Lenti-MPRA transduction, DNA/RNA library preparation and sequencing

Tonsillar GC B cells were isolated, cultured, and transduced with lenti-MPRA constructs as described above. In pilot analyses, we confirmed that the minP sequence was sufficient to drive GFP expression from the lenti-MPRA backbone in GC B cells, allowing us to detect both increases and decreases in transcriptional reporter activity. For the final MPRA library, on day 2 post-isolation tonsillar GC B cells from 2 donors (F3 and M6) were transduced with RD114-pseudotyped MSCV-BCL6-t2A-BCL2 to limit plasma cell differentiation, followed by transduction of 1.92 ×10^7^ GC B cells with GaLV-pseudotyped lentivirus MPRA library, with 3 independent technical replicate transductions performed per donor (6 replicates total). Cells were collected 3 days post-transduction for joint DNA and RNA isolation with Quick-DNA/RNA Miniprep Plus Kit (Zymo Research). Genomic DNA and cDNA libraries were constructed based on Gordon et al. (*87*) with custom primer sequences (MPRA_PCR1_XX_FW and MPRA_PCR1_mix_RV for PCR1; MPRA_PCR2_FW and MPRA_PCR2_RV for PCR2; table S13B) used for amplification. All libraries were pooled and sequenced on the Illumina NovaSeq X Plus with 150 bp paired-end reads to > 50 M reads per sample.

### Lenti-MPRA data processing and analysis

The resulting fastq files were trimmed, quality-filtered (q>20) and demultiplexed with cutadapt. Barcode and UMI sequences were extracted from each read, and UMI counts for each barcode were quantified with custom scripts. For each replicate, barcodes with ≥3 UMI counts in both RNA and DNA libraries were retained. These barcodes and previously obtained barcode-CRE associations were used to filter for CREs with ≥20 barcodes. For each CRE, RNA (CRE_RNA_) and DNA (CRE_DNA_) counts were determined as sum of the UMI counts for all assigned barcodes to that CRE in RNA and DNA libraries respectively and then normalised to the total number of UMI counts from all barcodes for RNA and DNA libraries respectively. Transcriptional efficiency (TE) was quantified as CRE_RNA_/CRE_DNA_ and then normalised to the median TE across all CREs in that replicate. Normalised TE values for REF- and ALT-containing sequences were compared using a Student’s t-test, and allele-specific fold differences were quantified as median(TE_ALT_)/ median(TE_REF_) (table S10).

### Prediction of allele-specific effects on transcription factor binding sites

The predicted effect of non-coding autoimmune risk variant alleles on transcription factor binding sites were assessed using motifbreakR (*88*). Position weight matrices (PWMs) from Jaspar (*89*) were matched to the hg38 genome using a threshold of *p* < 0.0001, and allelic effects on each PWM were quantified via the default sum of log-probabilities algorithm, specifying a background probability of 0.3 and 0.2 for A/T and G/C, respectively (table S11).

### Transcription factor binding at non-coding autoimmune risk loci

Human B cell REL ChIP-seq (*90*) was aligned to GRCh38 with bowtie2 (v2.4.4; *64*) before retaining uniquely mapped reads and duplicate read removal with samtools (v1.16.1; *65*), peakcalling with MACS2 (p <0.05 and FE >3; *66*) and bigwig files generated with deepTools (v3.5.1; *68*). Transcription factor ChIP-seq datasets for lymphoblastoid GM12878 B cells for the following were downloaded from ENCODE: CEBPB (ENCSR681NOM), ETS1 (ENCSR000BKA), IKZF1 (ENCSR441VHN), IRF3 (ENCSR408JQO), IRF4 (ENCSR000BGY) (*91, 92*). Peakcalling was performed with MACS2 for CEBPB (FE>4), ETS1 (FE>6), IRF3 (FE4), and IRF4 (FE>6), while for IKZF1 the previously published peak set was used.

### Prime editing (PE7) reagents and pegRNA design

Prime editing of autoimmune risk variants was performed with the PE7 system (PEMAX with additional LA factor) delivered as *in vitro* transcribed (IVT) mRNAs and synthetic pegRNAs (*93*). IVT production of PE7-tLNGFR and dnMLH1 was performed using modified constructs: pGEMs_PE7_CD271_IVT and pGEMs_MLH1dn_GFP plasmid templates, derived from the parental vectors pGEMs_eGFP_IVaxT_Poly_A_148 (*94*). Modified constructs were obtained as a gift from Charlotte Slade Lab and linearised by digestion with SapI (NEB) prior to purification using DNA Clean & Concentrator kit (Zymo Research). DNA was precipitated overnight with 5M NaCl to a final concentration of 0.1M and 3 volumes of 100% ethanol at −80°C before centrifugation for 10 minutes at 16,000 × *g* at 4°C. DNA pellets were air dried for 30 min at room temperature and reconstituted in nuclease-free water before storage at −20°C. Linearised plasmids were used to perform IVT using HiScribe T7 mRNA Kit with CleanCap Reagent AG (NEB) according to the manufacturer’s protocol, except for the following modification: substitution of UTP with N1-Methylpseudouridine-5’-Triphosphate (pseudo-UTP) (Trilink Biotechnologies). Reactions were treated with DNase I (RNase-free) (NEB) and incubated for 15 minutes at 37°C. IVT RNA was purified using Monarch RNA Cleanup kit (NEB), quantified by NanoDrop and quality assessed using the RNA screen tape on the 4200 TapeStation (Agilent). IVT RNA was diluted to a final concentration of 2 µg/µl and stored at −80°C. PE7-optimal pegRNAs (*95*) (table S13C) were synthesised by Genscript with following modifications to improve editing efficiency: 2’-O-methyl modification at the first three nucleotides, phosphorothioate linkages between the first three and last three nucleotides, and UU*mU*mU*mUU encoded at the 3’ end (*93*).

### Prime editing of non-coding risk variants in primary human B cells

GC B cells were isolated from frozen tonsil mononuclear cells and cultured for 4 days on pre-plated YK6 feeder cells as described above. Cells were harvested and counted, before 1-3 x 10^6^ cells were electroporated with 2 µg PE7-tLNGFR mRNA, 2 µg dnMLH1 IVT mRNA and 50 pmol synthetic pegRNA (table S13C) using the P3 Primary Cell 4D-Nucleofector® X kit (Lonza Bioscience) and the EO-117 program on the Lonza 4D-Nucleofector®. Cells nucleofected with PE7-tLNGFR and dnMLH1 without synthetic pegRNA were used as negative controls. Cells were cultured for a further 4 days post-nucleofection before harvesting and FACS-based analysis.

### Quantile-based analysis of target gene expression by sequencing

Prime editing efficiency rates are often low and highly variable between different loci. To standardise our experimental pipeline and quantify the genetic effects of autoimmune risk variants on target gene expression, we implemented a quantile-based analysis of target gene expression by sequencing workflow inspired by recent studies (*24, 96*). For experiments measuring differential effects of targets expressed on the cell surface, cells nucleofected with prime editing components were stained with relevant antibodies: CD83-APC (HB15e, Biolegend), CD84-APC (CD84.1.21, Biolegend), TNFSF4(OX40L)/CD252-PE (11C3.1, Biolegend), SLAMF1/CD150-PE (A12 (7D4), Biolegend), SLAMF6/CD352-PE (NT-7, Biolegend) at a 1:50 dilution for 30 minutes on ice in the dark and washed three times with FACS buffer before sorting. For experiments measuring differential effects of *REL* transcript levels, cells nucleofected with prime editing components to introduce rs1432296 C>T were stained with PrimeFlow™ REL-Alexa Fluor 647 (Thermo Fisher Scientific) and the PrimeFlow™ RNA Assay kit (Thermo Fisher Scientific) following the manufacturer’s protocol with 3-10 x 10^6^ cells for fixation and permeabilization, and a maximum of 5 x 10^6^ cells for target probe hybridization. Cells were sorted into quartiles based on target fluorescence using the Aurora CS Cell sorter, BD FACS Aria FUVsion, BD FACS Aria ConFusion d’Fusion, with each bin comprising 20-25% of cells, prior to genomic DNA isolation with Quick-DNA Miniprep Plus or Quick-DNA Microprep Plus (Zymo Research) for antibody-labelled samples. For PrimeFlow™ experiments, samples were incubated with proteinase K at 56°C overnight followed by 90°C for 2 hours to reverse crosslinking and DNA isolated with QIAamp DNA FFPE Tissue Kit (Qiagen).

To quantify edited (risk) variant allele frequencies in different target expression quartiles, >50 ng gDNA was amplified with NEBNext Ultra II Q5 Master Mix (NEB) and 0.5 µM PE_rsX_FW and PE_rsX_RV primers (table S13B) with the following conditions: 98°C for 30 s, (98°C for 10 s, 63∼66°C for 30s, 72°C for 30s) × 12 cycles, 72°C for 5 min. PCR products were purified using 1X AMPure XP beads (Beckman Coulter) and prior to amplification and addition of indexing barcodes with NEBNext Ultra II Q5 Master Mix and 0.5 µM PE_PCR2_FWX and PE_PCR2_RVX primers (table S13B) with the following conditions: 98°C for 30 s, (98°C for 10 s, 65°C for 30s, 72°C for 30s) × 24 cycles, 72°C for 5 min. Indexed libraries were purified with 0.8X AMPure XP beads, quantified with Qubit™ dsDNA HS Assay Kit (Invitrogen) and pooled together at equimolar ratios before addition of Oxford Nanopore adapters as a single library pool with Ligation Sequencing Kit V14 (ONT, SQK-LSK114) and sequencing on the Oxford Nanopore PromethION 2 Solo. Oxford Nanopore pod5 files underwent basecalling with the super-accurate model (dna_r10.4.1_e8.2_400bps_sup@v5.0.0) of dorado v0.7.3 (Oxford Nanopore Technologies) and resulting fastq files were demultiplexed and quality-filtered (q>20) using cutadapt. The nucleotide at the edited position was determined by trimming 5’and 3’flanking genomic sequences with cutadapt, and count tables with were generated for each sample using custom scripts. The frequency of each nucleotide at the edited position was determined as the number of counts for this nucleotide divided by the total count, with further normalisation to the mean frequency of this nucleotide across 4 quantiles for a given donor.

## Acknowledgements

The authors thank Johannes Wichmann, Daniela Zalcenstein, Casey Anttila, Ling Ling, Ruvimbo Mishi, and Rory Bowden of WEHI Advanced Genomics Facility and all team members of the WEHI Flow Cytometry Facility for their support and assistance in this work. We thank Sarahi Mendoza Rivera and Miles Horton for helpful discussions about prime editing strategies. We thank the King Lab consumer group members Eliza Metcalfe and L. E. Ohlman for their support. Some figures were made with the assistance of BioRender. We thank all the parents, children, and volunteers who participated in this study.

## Author contributions

V.A.K. designed, executed and analysed nearly all experiments. J.W.H.C. and J.L. designed and executed prime editing experiments. D.V. and H.W.K. contributed to bioinformatic analysis. M.N., E.L., K.D., S.S. and V.L.B. provided access to tonsillectomy patient biospecimens. L.G. assisted with tissue processing. G.S. and D.J.H. provided key reagents and critical advice on viral delivery of CRISPR-SAM system into primary B cell cultures. N.D. and J.C.H. performed micro-Capture-C experiments and data analysis, for which J.O.J.D. supervised. E.B.S. provided key reagents and advice for in vitro transcription and prime editing protocols. V.A.K. and H.W.K. wrote the manuscript with input from all authors. The project was conceived and supervised by H.W.K.

## Data, code, and materials availability

Processed single-cell gene expression, cell surface protein and sgRNA count matrices are available at Zenodo (10.5281/zenodo.18828272). All other raw and processed sequencing datasets are currently awaiting upload to online data repositories and will be made available upon request.

## Supplementary Materials

Table S1. Curated set of 5655 fine-mapped autoimmune risk SNPs.

Table S2. Prioritised set of 818 autoimmune SNPs accessible in B cell open chromatin.

Table S3. SCANDAL sgRNA library targeting 763 autoimmune risk loci.

Table S4. Summary of SCANDAL loci with mapped *cis*-regulatory gene targets.

Table S5. Full result table from sceptre differential gene expression analysis.

Table S6. Full result table from sceptre differential protein expression analysis.

Table S7. 5’ TSS RNA expression at SCANDAL CREs before and after CRISPR activation.

Table S8. Result table from sceptre differential CRE expression analysis at SCANDAL CREs before and after CRISPR activation.

Table S9. Result table from testing for intergenic CRE-CRE coregulation pairs with sceptre.

Table S10. MPRA result summary.

Table S11. motifbreakR result summary.

Table S12. c-REL-bound SCANDAL loci and *cis*-regulatory gene targets.

Table S13. Sequences of used sgRNAs, pegRNAs, and primers.

Table S14. Dictionary of autoimmune traits.

Table S15. MPRA sequences.

**Fig. S1.**
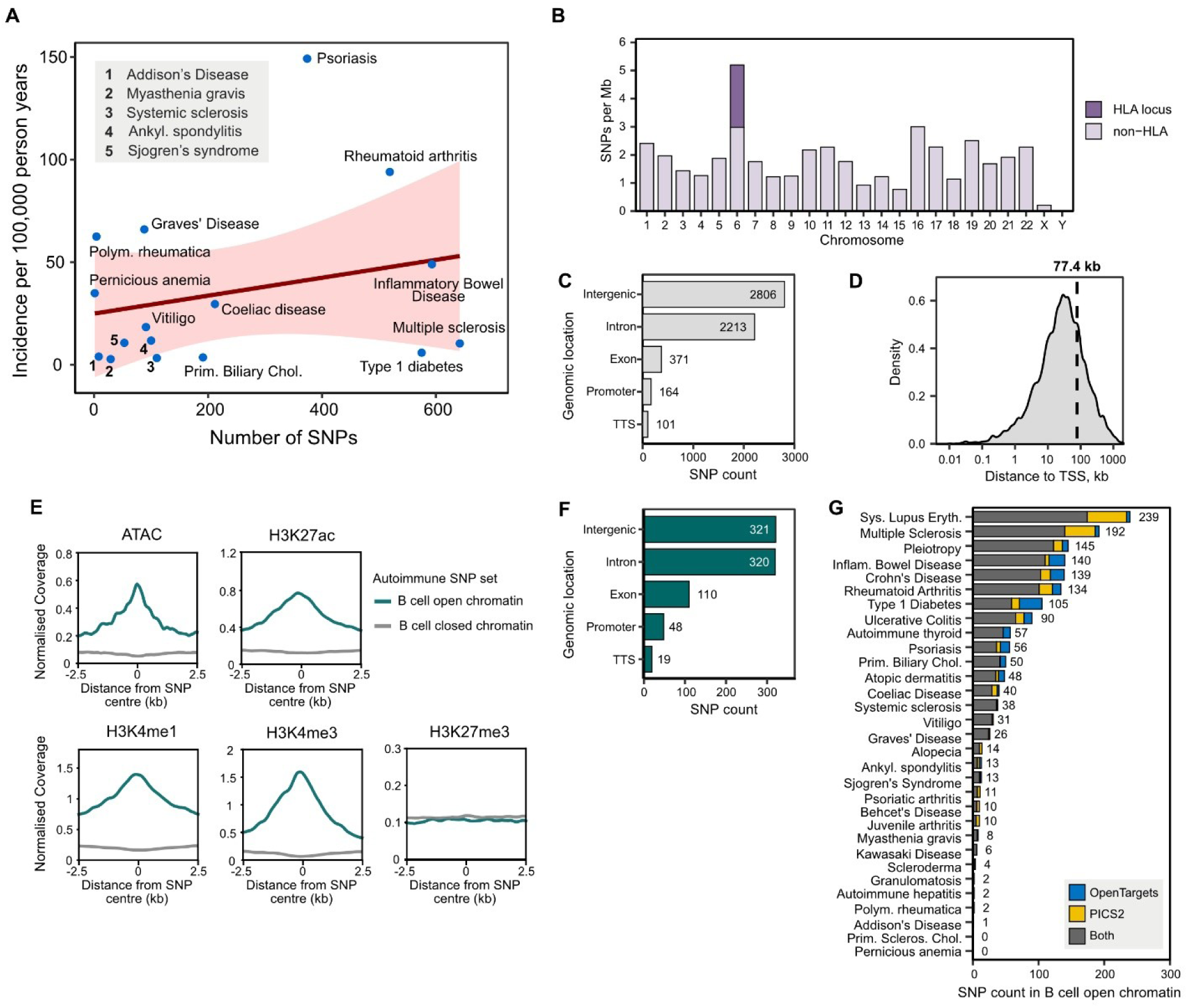
Features of pan-autoimmune disease GWAS risk variant set. (**A**) Comparison between autoimmune trait frequency and amount of GWAS data. (**B**) SNP distribution across chromosomes. (**C**) Number of SNPs in different genomic locations. TTS = transcription termination site. (**D**) Distance from SNPs to their nearest TSS. Dashed line shows the mean distance. (**E**) Comparisons between GC B cell chromatin accessibility (ATAC) and histone modifications between SNPs that fall within B cell open chromatin ATAC-seq peaks (818 SNPs) and those that do not overlap open chromatin (4837 SNPs). (**F**) Genomic locations of 818 SNPs within B cell open chromatin. (**G**) Trait frequency for 818 autoimmune risk SNPs within B cell open chromatin from PICS2 and OpenTargets Genetics.

**Fig. S2.**
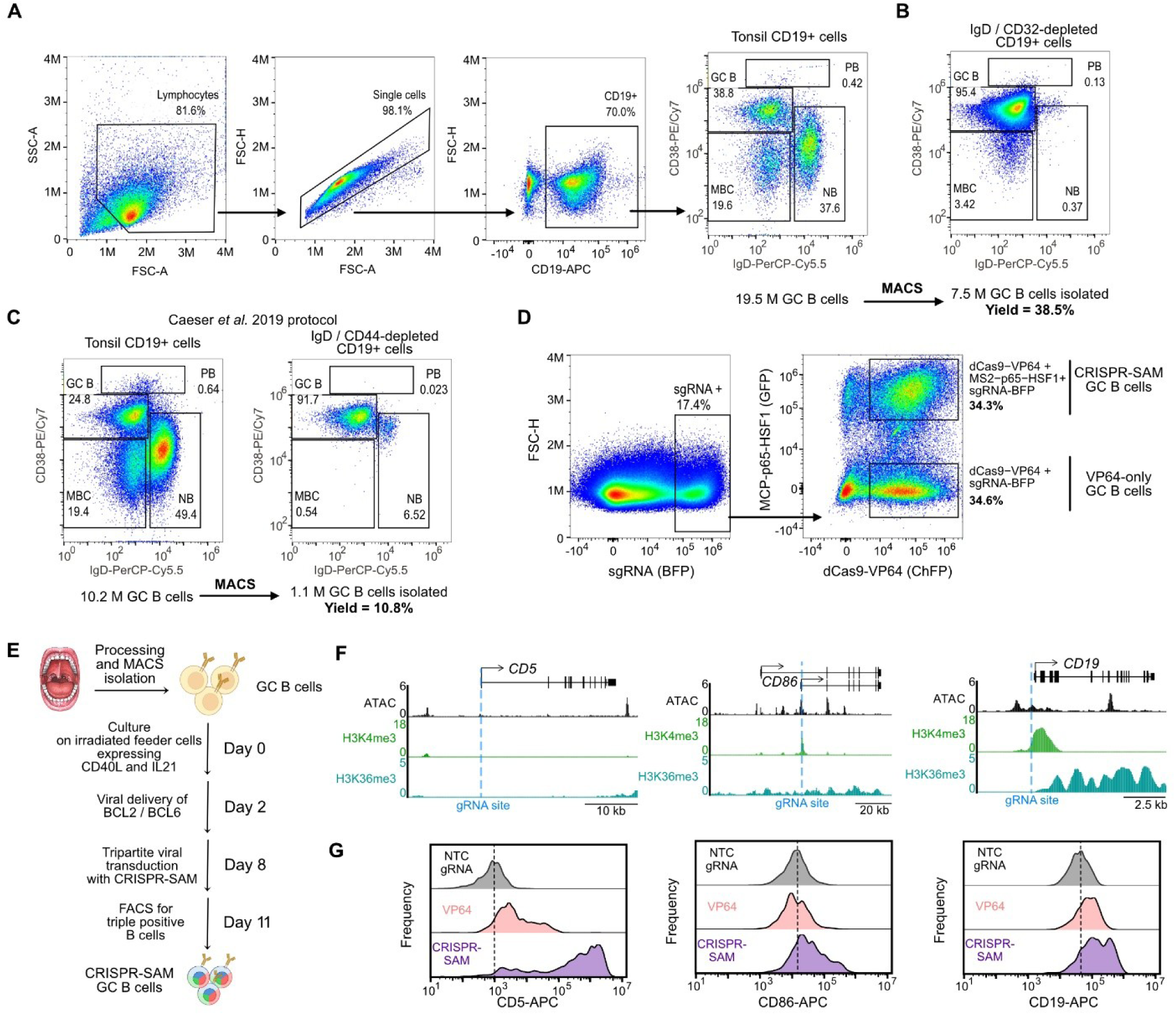
Isolation and CRISPR activation in human tonsillar GC B cells. (**A**) FACS gating strategy and frequencies of different tonsillar CD19+ B cell subsets (NB = Naïve B cells, GC B = germinal centre B cells, MBC = memory B cells, PB = plasmablasts). (**B**) Enrichment and yield of GC B cells following IgD- and CD32-depletion. (**C**) Enrichment and yield of GC B cells following IgD- and CD44-depletion as published by Caeser *et al*. (**D**) FACS sorting strategy for triple (CRISPR-SAM + sgRNA) and double (dCas9-VP64 + sgRNA) positive GC B cells 3 days post-transduction. (**E**) Schematic and timeline for CRISPRa experiment with BCL2/BCL6 over-expression step. (**F**) Genome snapshots of three different promoters (*CD5, CD86, CD19*) targeted for CRISPR activation. (**G**) Representative FACS plots from *CD5, CD86, CD19* CRISPR activation experiments. Dashed line shows the median MFI in non-targeting controls.

**Fig. S3.**
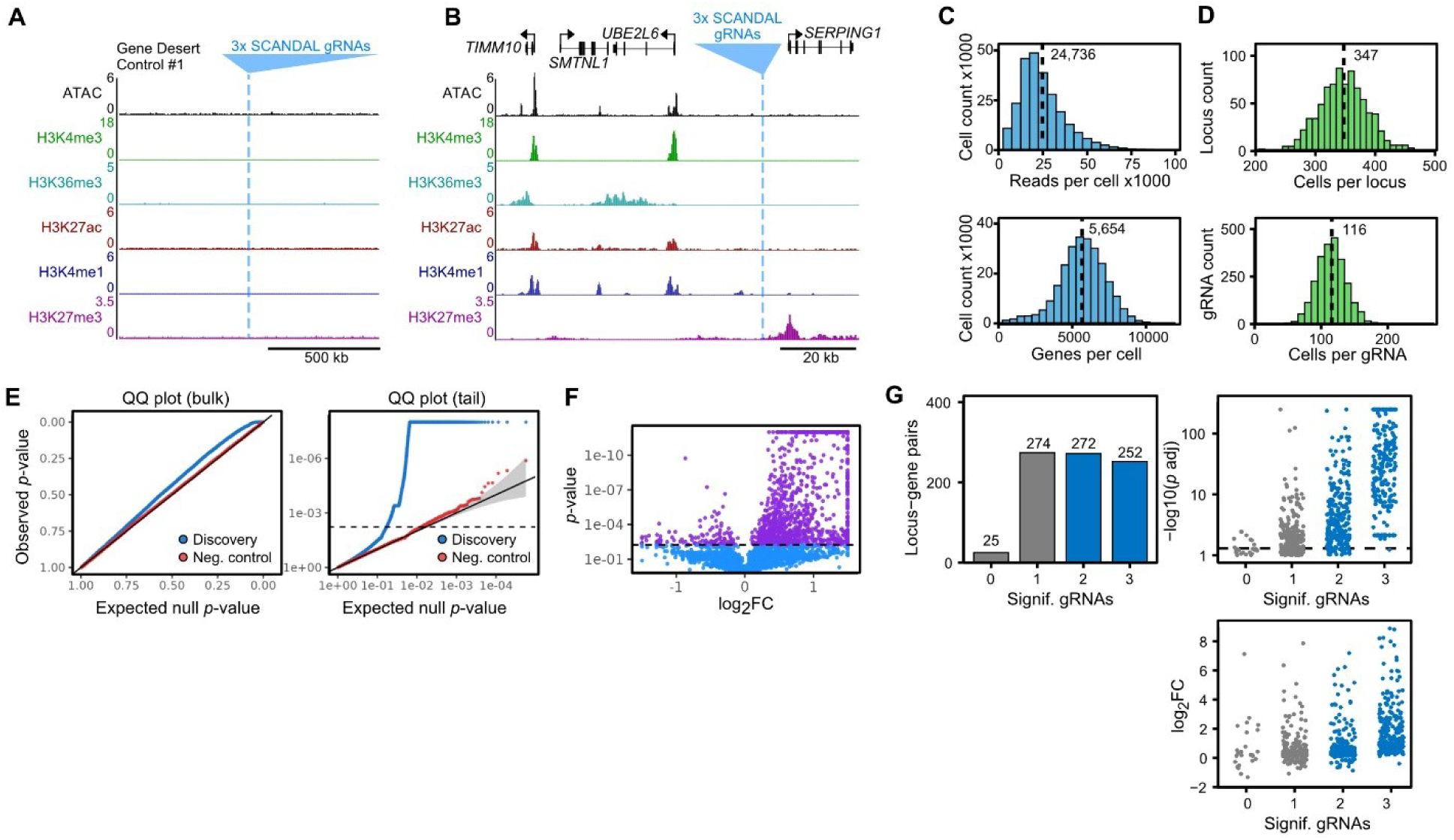
Single-cell CRISPR activation screen design and statistical testing with SCEPTRE. (**A**) Genome snapshots of representative negative gene desert control in SCANDAL. (**B**) As in (A), but for distal control (closed chromatin in proximity to TSS). (**C**) Summary statistics from SCANDAL sequencing and analysis, including number of reads per cell and number of detected genes per cell. (**D**) SCANDAL library representation, including number of cells per locus (three sgRNAs) and per sgRNA. (**E**) Calibration check analysis with SCEPTRE, including differences in *p*-values for negative controls (NTC sgRNAs) and discovery pairs (gRNAs targeting risk and control loci). (**F**) Discovery volcano plot from SCEPTRE analysis. Each dot represents a locus-gene pair. All pairs that were called as significant are shown in purple. (**G**) Properties of significant locus-gene pairs identified by SCEPTRE from a pooled analysis of all three sgRNAs per locus. Loci are grouped according to the number of individual sgRNAs that returned a significant locus-gene effect. 0 sgRNAs denote locus-gene pairs that were only significant in the pooled analysis.

**Fig. S4.**
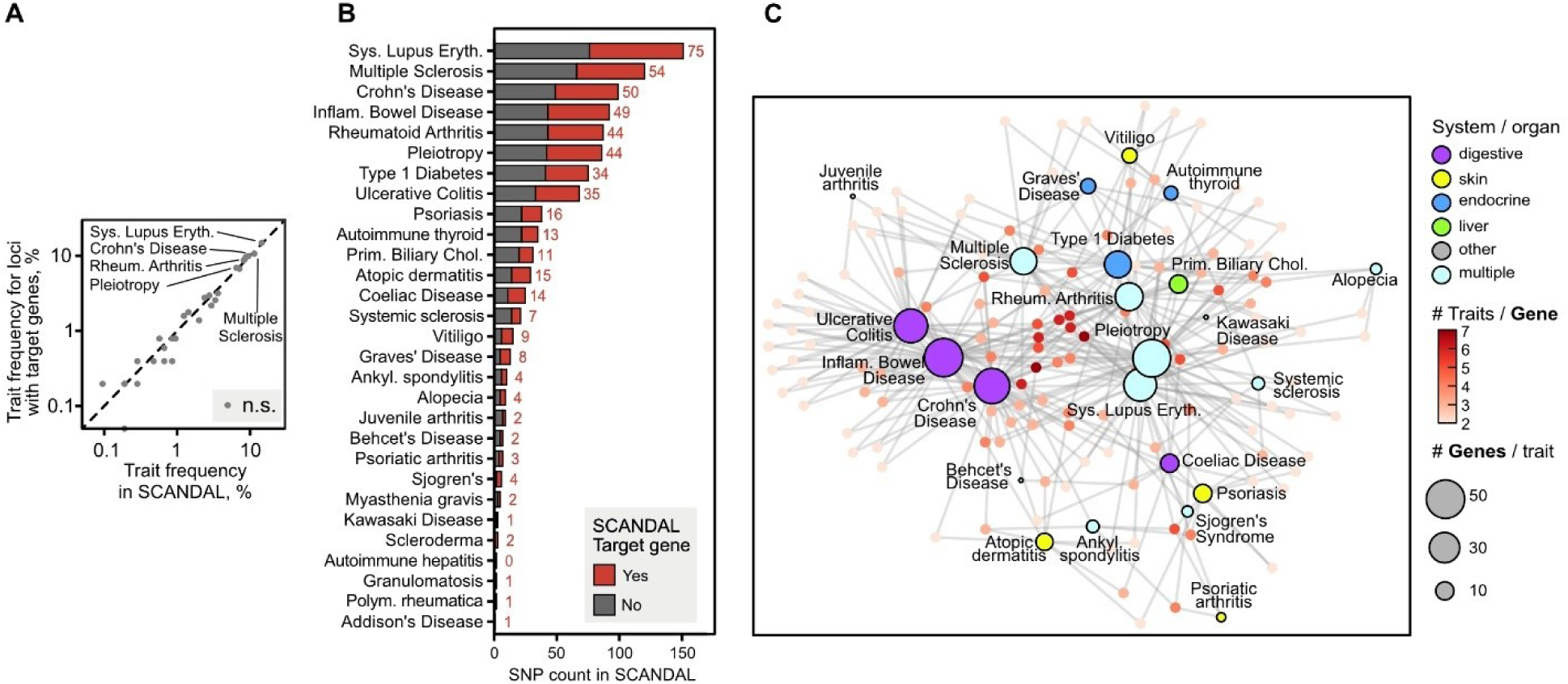
Trait analysis from locus-gene pairs identified by SCANDAL. (**A**) Trait enrichment for SCANDAL risk loci with mapped gene targets compared to complete SCANDAL risk loci, as determined by Fisher exact test. n.s. denotes not significant (*p* > 0.05). (**B**) Number of SCANDAL risk loci with identified *cis*-regulatory target genes for each GWAS autoimmune trait. (**C**) Shared trait associations for multi-trait SCANDAL target genes visualised as a force-directed graph.

**Fig. S5.**
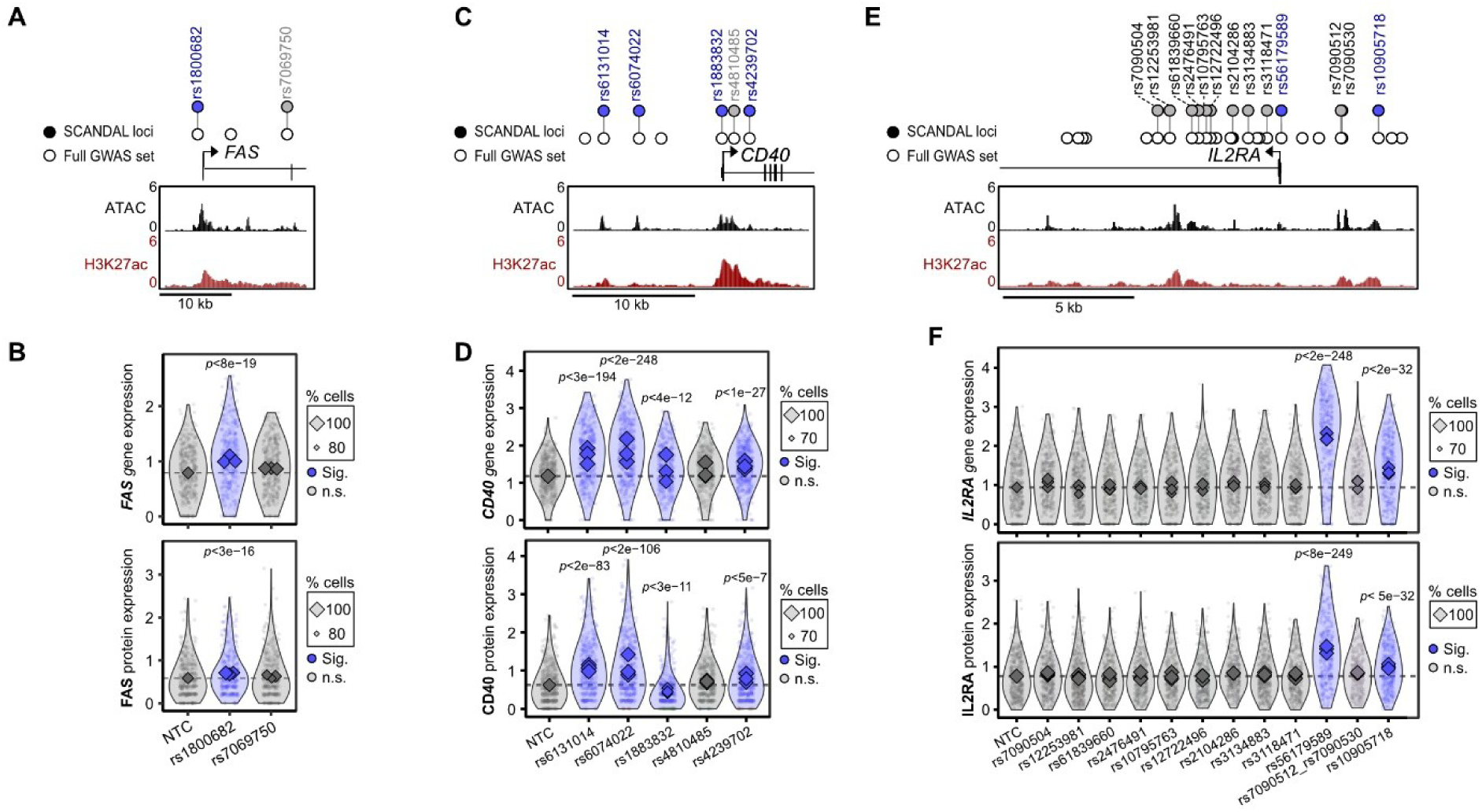
Analysis of matched gene and protein expression following CRISPR-SAM activation of risk loci in SCANDAL. (**A**) Genome snapshot of CRISPR-SAM targeted risk loci in proximity to *FAS*. SCANDAL-tested loci are shown in filled circles while complete autoimmune GWAS SNP set are shown as white circles. (**B**) Gene and protein expression for *FAS*/FAS following CRISPR-SAM activation of SCANDAL loci. Circular points reflect single cell expression values, while diamonds indicate mean expression values for all cells containing individual sgRNAs where size corresponds to percentage of cells in which the gene was detected. Benjamini-Hochberg-adjusted SCEPTRE *p*-values are shown. (**C**) Genome snapshot of CRISPR-SAM targeted risk loci in proximity to *CD40*. (**D**) Gene and protein expression for *CD40*/CD40 following CRISPR-SAM activation of SCANDAL loci. (**E**) Genome snapshot of CRISPR-SAM targeted risk loci in proximity to *IL2RA*. (**F**) Gene and protein expression for *IL2RA*/IL2RA following CRISPR-SAM activation of SCANDAL loci.

**Fig. S6.**
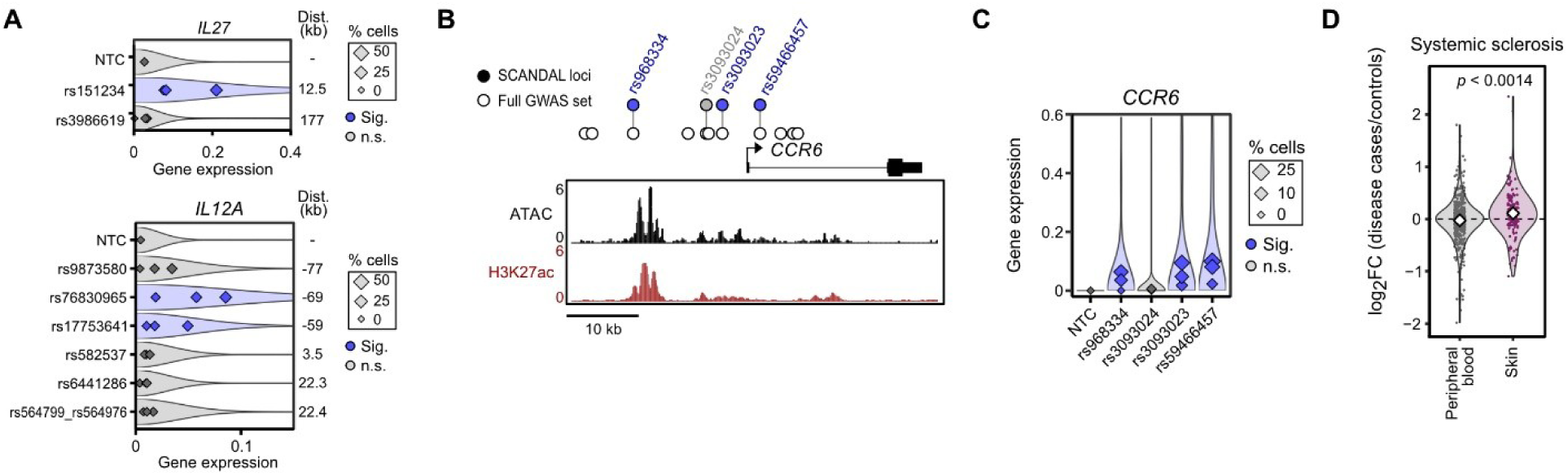
Examples of lowly expressed SCANDAL target genes and disease-relevant expression. (**A**) Perturbation effects of SCANDAL target genes that are lowly expressed at baseline. Significant effects are shown in blue. (**B**) Genome snapshot of CRISPR-SAM targeted *CCR6* locus. (**C**) Single-cell expression perturbation effects of *CCR6*. Significant effects per locus are shown in blue. (**D**) Fold change in target gene expression between disease (Systemic sclerosis) and control cases in peripheral blood and disease-relevant tissue (skin). *p*-value from Student’s t-test is shown.

**Fig. S7.**
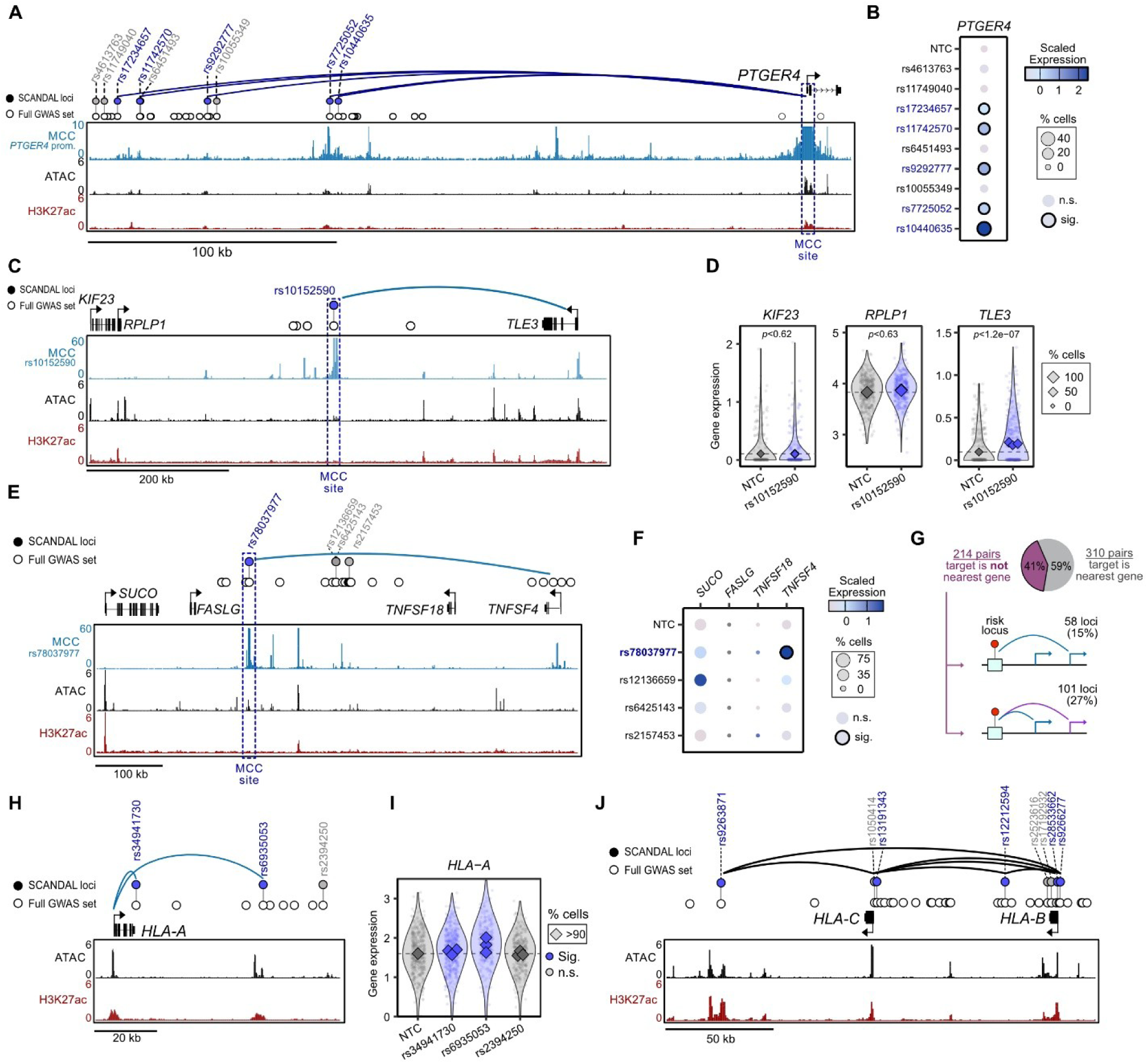
Different patterns of gene regulation by distal autoimmune risk loci. (**A**) Genome snapshot of CRISPR-SAM targeting and Micro-Capture-C (MCC) for SCANDAL loci upstream of *PTGER4*. (**B**) Scaled expression (z-scores) of *PTGER4* for cells containing sgRNAs targeting SCANDAL loci or non-targeting control sgRNAs (NTC). (**C**) Genome snapshot of CRISPR-SAM targeting and Micro-Capture-C (MCC) for rs10152590, including SCANDAL *cis*-regulatory effect on target gene *TLE3* highlighted as curved line. (**D**) Expression of *KIF23*, *RPLP1* and *TLE3* near rs10152590 for cells containing rs10152590-targeting or non-targeting control sgRNAs. Circular points reflect single cell expression values, while diamonds indicate mean expression values for all cells containing individual sgRNAs where size corresponds to percentage of cells in which the gene was detected. *p* denotes Benjamini-Hochberg-adjusted SCEPTRE *p*-value. (**E**) Genome snapshot of CRISPR-SAM targeting and Micro-Capture-C (MCC) for rs78037977, including SCANDAL *cis*-regulatory effect on target gene *CCND3* highlighted as curved line. (**F**) Scaled expression (z-scores) of *TNFSF4* and nearby genes for cells containing sgRNAs targeting SCANDAL loci or non-targeting control sgRNAs (NTC). (**G**) Frequency that gene target for a risk locus is the nearest gene or not, and frequency of gene skipping or multiple gene target SCANDAL categories. (**H**) Genome snapshot of CRISPR-SAM targeting for *HLA-A*-regulating loci, including SCANDAL *cis*-regulatory effects on target gene *HLA-A* highlighted as curved line. (**I**) Expression levels of *HLA-A* for cells containing loci-targeting or non-targeting control sgRNAs. (**J**) Genomic context of SCANDAL risk loci and SNPs at *HLA-C* and *HLA-B*.

**Fig. S8.**
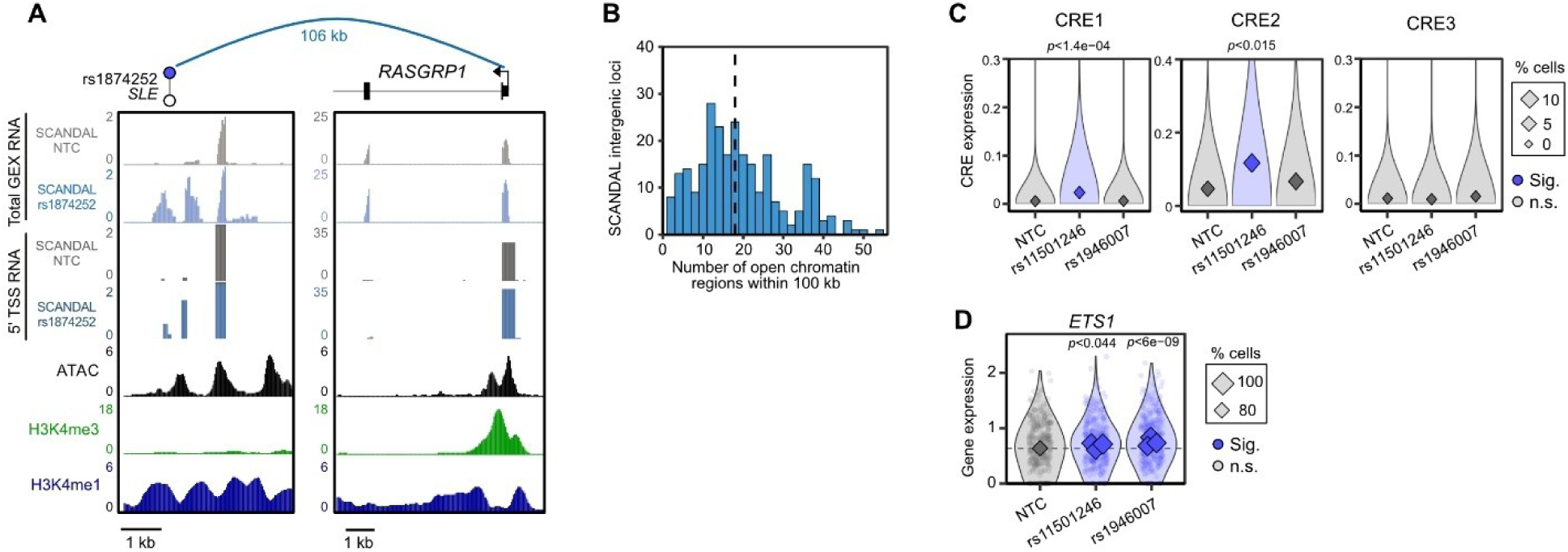
Transcriptional activity and proximity of nearby CREs at SCANDAL loci. (**A**) Genome snapshots of CRISPR-SAM targeting perturbation effects for rs1874252 and the TSS of *cis*-regulatory target gene *RASGRP1* (106 kb upstream), including tCRE activity (total GEX RNA and 5’ TSS RNA quantified by SCAFE) for cells containing non-targeting control sgRNAs and cells containing sgRNAs targeting rs1874252. (**B**) Number of open chromatin regions within 100 kb of SCANDAL intergenic loci. Dashed line shows the median. (**C**) tCRE activity (total GEX RNA) of CREs for cells containing non-targeting control sgRNAs and rs11501246-targeting or rs1946007-targeting sgRNAs. Diamonds indicate mean expression values for all cells containing sgRNAs where size corresponds to percentage of cells in which the tCRE was detected. *p* denotes Benjamini-Hochberg-adjusted SCEPTRE p-value. (**D**) Expression of *ETS1* for SCANDAL loci described in (C). *p* denotes Benjamini-Hochberg-adjusted SCEPTRE *p*-value.

**Fig. S9.**
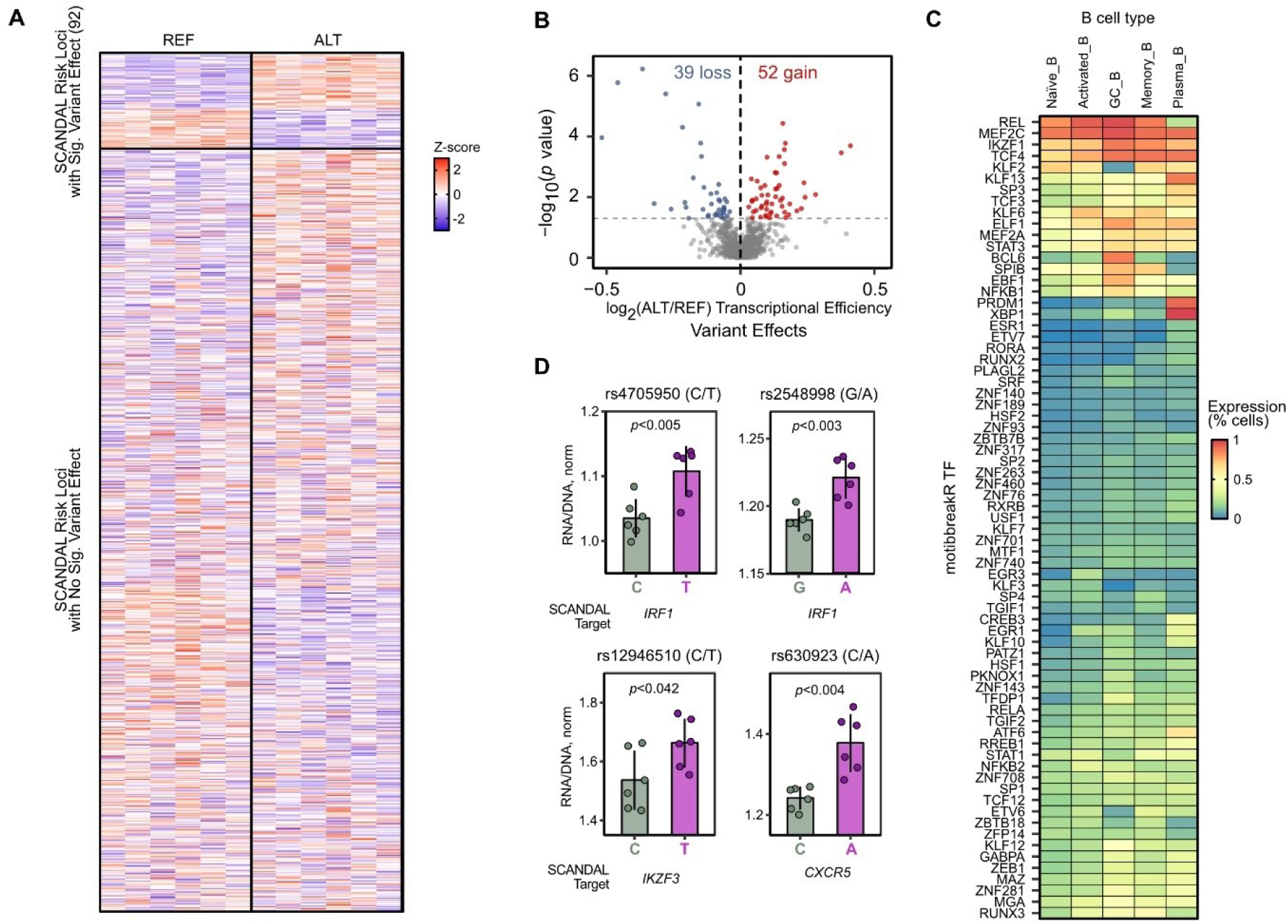
LentiMPRA to quantify variant-specific effects at SCANDAL loci and expression of transcription factors identified from motifbreakR. (**A**) Reporter gene expression (z-score) for reference (REF) and alternate (ALT) alleles for 818 SCANDAL loci. (**B**) Differential variant-specific effects from lentiMPRA in GC B cells. Horizontal dashed line corresponds to p-value of 0.05. (**C**) Transcription factors (TF) with strong motifbreakR prediction of motif disruption by SNP and their expression in B cell types. (**D**) Normalised transcriptional efficiency (RNA/DNA, norm) for reference (grey) or alternate (purple) alleles at SCANDAL autoimmune risk loci encoding rs4705950 (*IRF1*), rs2548998 (*IRF1*), rs12946510 (*IKZF1*) and rs630923 (*CXCR5*). Genes in brackets denote experimentally-derived SCANDAL *cis*-regulatory target gene, and *p* denotes results from paired sample t-test (n = 6, ±SD).

**Fig. S10.**
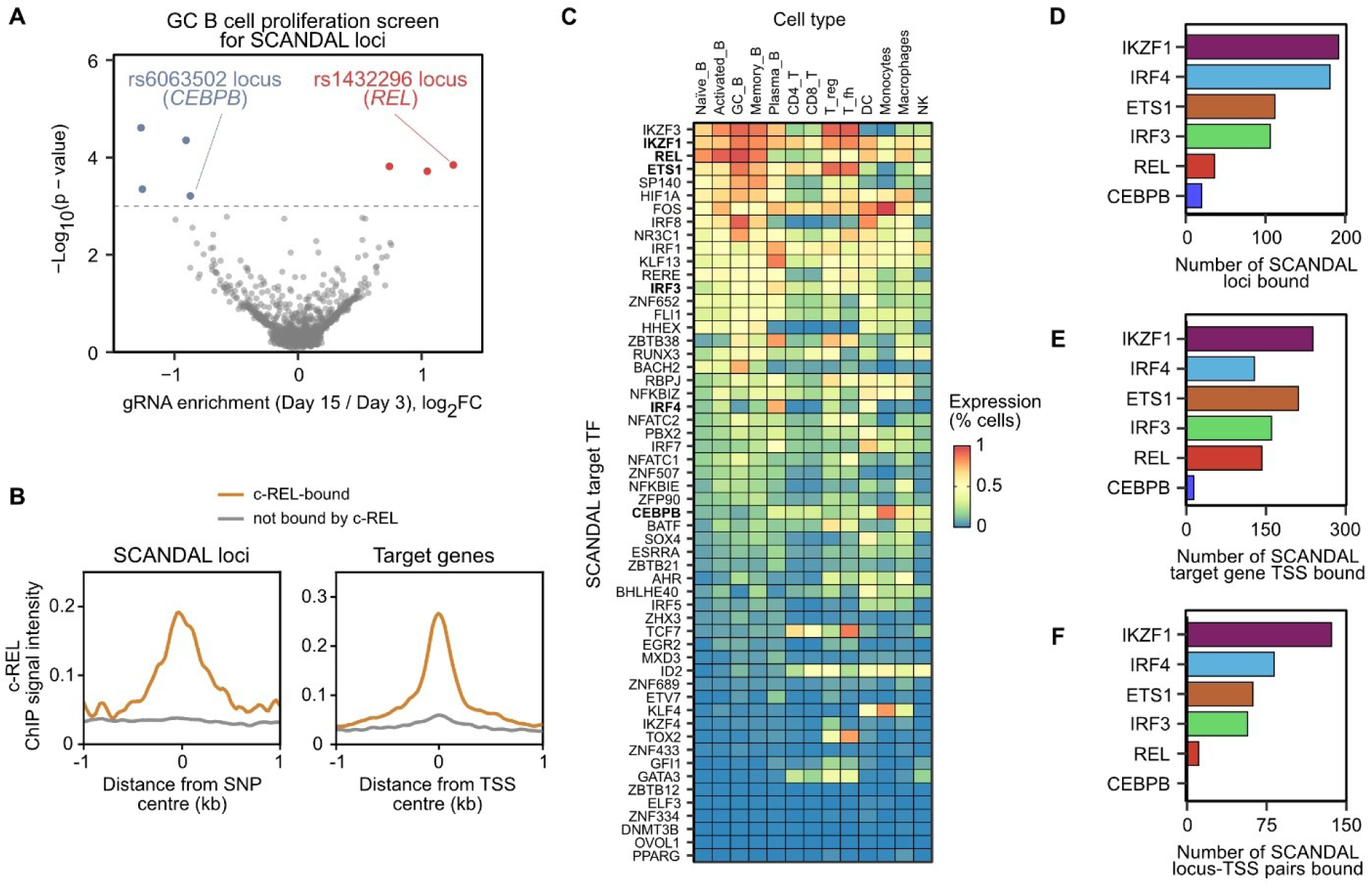
Transcription factors identified as SCANDAL targets extensively bind at other autoimmune risk loci and their target genes. (**A**) Results from bulk CRISPRa proliferation screen showing SCANDAL loci whose activation either increased or decreased the proliferation of GC B cells. (**B**) c-REL ChIP signal at autoimmune risk loci and 362 target genes from Naïve B cells. (**C**) 56 transcription factor (TF) genes identified as SCANDAL targets and their expression in B cell types. (**D-F**) Binding of SCANDAL target TFs at (**D**) SCANDAL loci, (**E**) TSS of 362 SCANDAL target gene TSS, (**F**) both locus and TSS of its target gene.

